# Isotropic, aberration-corrected light sheet microscopy for rapid high-resolution imaging of cleared tissue

**DOI:** 10.1101/2025.02.21.639411

**Authors:** Mostafa Aakhte, Gesine F. Müller, Lennart Roos, Joe Li, Torben Göpel, Kurt R. Weiss, Aleyna M. Diniz, Jan Wenzel, Markus Schwaninger, Tobias Moser, Jan Huisken

## Abstract

Light sheet microscopy is the ideal technique for multiscale imaging of large and cleared tissues, and it is desirable to achieve the highest possible isotropic resolution across the entire sample. However, isotropic resolution for a centimeter-sized sample has only been achieved with slow and often aberrated, axially scanned light sheets, resulting in a low resolution of several micrometers. Here, we introduce a compact, high-speed light sheet fluorescence microscope with isotropic sub-micron resolution optimized for cleared tissue. We introduce three major opto-mechanical innovations using off-the-shelf optics to achieve an isotropic resolution of 850 nm across samples up to 1 cm^3^ and refractive indices ranging from 1.33 to 1.56, using mechanical tiling with a field of view of 800 µm × 800 µm. We show that combining an air objective and a meniscus lens achieves an axially swept light sheet with sub-micron diffraction-limited resolution and aberration correction. The effective field of view is increased 2-fold by correcting the field curvature of the light sheet with a concave mirror in the remote focusing unit. Furthermore, the imaging speed is enhanced 10-fold by adapting the light sheet’s motion with a closed-loop feedback, reaching 100 frames per second while maintaining isotropic resolution across the large field of view. Finally, we showcase the performance of our light sheet system for imaging from subcellular up to centimeter scale in cleared zebrafish, mouse cochlea, and mouse brain using various clearing methods.

## Introduction

Light sheet fluorescence microscopy (LSFM) has become a key instrument in modern biology with its confined excitation and its capability to image live, intact biological samples fast and gently^1–4^. In LSFM, optical sectioning is achieved by two orthogonal optical arms^5^: one projects a laser light sheet into the sample, illuminating a thin section, and the other records the emitted fluorescence light perpendicularly with a camera. This configuration enables simultaneous collection of emitted light across the entire field of view (FOV), surpassing sequential point-scanning methods in photon efficiency and speed. Localized fluorescence excitation at the chosen plane significantly reduces photobleaching, which is critical for in vivo imaging, but LSFM is also an effective method for imaging large and cleared tissues6–10. It enables scientists to obtain high-resolution images across the entire intact sample, crucial for subsequent reconstructions and quantifications.

The light sheet properties, such as its thickness, effective length (Rayleigh length), and intensity distribution in all directions greatly influence image quality. The thickness of the light sheet primarily determines the axial resolution. In contrast, the lateral resolution is determined by the detection objective lens’ numerical aperture (NA), magnification, and camera pixel size^5,11^. Consequently, the axial resolution can be enhanced by creating thinner light sheets. However, a thinner light sheet becomes shorter, not filling the entire FOV anymore. Quantitative multiscale analysis requires the imaging setup to have an isotropic resolution, where the axial and lateral resolutions are the same across the entire FOV, and, consequently, the illumination and detection NA need to identical. Isotropic resolution across a large FOV can elegantly be achieved by rapidly moving a thin light sheet across the FOV using a remote-focusing optical arm, a technique known as axially swept light sheet microscopy (ASLM)^12,13^. To record only the thinnest part of the light sheet across the FOV, the camera is oriented such that the rolling shutter direction aligns with the light sheet’s propagation direction (**Supplementary Movie 1**). The light sheet is swept across the FOV once during the exposure time of the sCMOS (scientific complementary metal–oxide–semiconductor) camera, with its waist location synchronized to the position of the camera’s rolling shutter.

In ASLM, the light sheet can be swept with electrically tunable lenses (ETLs), piezo actuators, or voice coil actuators. The concept was initially introduced using a piezo actuator for imaging small live samples, providing an isotropic resolution of 370 nm over a 50 μm FOV at an imaging rate of 6 Hz^12^. ASLM was also adapted to larger cleared samples, as demonstrated in the MesoSPIM platform, which achieved a FOV of 13.29 mm with an isotropic resolution of 6.5 μm^14^. The subsequent benchtop MesoSPIM variant offered an improved resolution, albeit non-isotropic, at 1.5 μm laterally and 3.3 μm axially, with frame rates of 4.5 Hz and 6.5 Hz using ETLs^15^. Another setup achieved isotropic resolution up to 380 nm, covering a FOV of 310 × 310 μm^2^ and using mechanical tiling to image large samples^13^. This setup employed a voice coil actuator, which provided higher stroke movement in continuous motion, increasing the frame rate to 10 Hz. However, the setups, so far, have not reached the maximum frame rate of the sCMOS camera. The speed, however, is crucial to keep acquisition times within reasonable limits, especially in large tissues where many tiles need to be acquired at high resolution to reconstruct the entire sample.

To support a wide range of applications, the microscope should be compatible with expansion microscopy^16,17^ and many tissue-clearing protocols such as 3DISCO, iDISCO, EZ Clear, etc.^18–20^ and their broad range of refractive indices (RIs), from RI=1.33 (water) to RI=1.56 (Ethyl cinnamate; ECi). A straightforward approach to image samples with various RIs uses air objective lenses with long working distances^21,22^. Oily and toxic solvents are then enclosed in a chamber with glass windows, and the air lenses can easily be exchanged without touching the solvent. However, in such a configuration, the lenses induce substantial spherical, chromatic aberrations and field curvature, preventing reaching the diffraction limit. Spherical aberrations can be mitigated by introducing a meniscus lens instead of the flat glass window between the air objective lens and immersion media^23^. An open-top light sheet microscope^24,25^ used a customized meniscus lens to achieve a near-diffraction-limited light sheet. However, a low NA light sheet in combination with a high NA detection lens limited resolution to 1.1 μm laterally and 2 μm axially with an effective FOV of 780 μm x 375 μm because of aberrations at the edge of the detection arm.

Field curvature is also a common aberration, limiting the FOV and affecting both the detection and illumination arms. In the detection arm, field curvature results in blur at the edges of the image. On the illumination side, the waist of the light sheet is bent in the presence of field curvature, which is particularly pronounced for high NA. This effect leads to a desynchronization of the waist movement at the FOV edges in ASLM and reduces the usable FOV to a small area where the waist is straight. The field curvature can be addressed by reducing the FOV or designing an expensive custom objective lens^26^. One can also select lenses with minimum field curvature by measuring their field curvature^15,27^. Further, the existing field curvature can be adapted by adjusting the light sheet geometry and using a curved beam^28^. Although all the above-mentioned efforts use at least one air objective lens with minimized aberrations, each approach falls short of achieving isotropic sub-micron resolution over the largest possible FOV.

To overcome all these issues, we have developed a compact, field curvature-corrected light sheet microscope tailored for imaging cleared tissues. The microscope offers an isotropic resolution of 850 nm in media with RIs ranging from 1.33 to 1.56 at a speed of 100 frames per second to image samples up to 1 cm^3^ in size. Our innovative design combines a multi-immersion lens, an air objective lens, a meniscus lens, and a concave mirror within an ASLM framework. We have doubled the FOV by using a concave mirror to correct the field curvature of the light sheet while maintaining isotropic resolution. Further, we have greatly improved the imaging speed by a factor of 10, compared to the fastest previous microscopes, by precisely controlling the ASLM unit at high frequencies with an optical closed loop. We benchmarked the system performance by imaging various biological samples, including cleared mouse brain, mouse cochlea, and zebrafish.

## Results

We aimed to achieve an isotropic resolution <1 µm to image sub-micron structures in large, cleared samples in various media. Hence, the detection objective lens needed to have an NA of ≥0.4. Commercially available options included a multi-immersion lens (16x, NA 0.4, working distance ca. 12 mm, ASI), which achieves optimal sampling when paired with a 200 mm tube lens and an sCMOS camera with a typical pixel pitch of 6.5 μm (**Supplementary Note 1 and Supplementary Fig. 1**). This lens is already corrected for chromatic aberrations with constant working distance for different RIs, compatible with various media without the need to realign the lens when changing the medium, making it the ideal choice for the detection lens in our setup.

Conversely, an air objective lens with matching NA=0.4 served as an ideal illumination lens. We placed it outside the interchangeable sample chamber and its toxic solvents. The air lens can be freely moved during alignment without requiring any sealing. The flexibility of the air illumination objective lens also enabled us to attach the detection lens in a fixed position, sealed to the chamber with a simple O-ring, and its focal plane in the chamber center aligned to the illumination’s optical axis. The illumination lens’ large working distance (20 mm in air) is beneficial for imaging large samples of a centimeter or more. The chosen Mitutoyo objective lens (20X plan apochromat) has an NA of 0.42, which closely matches the detection lens’ NA, a prerequisite for achieving isotropic resolution. In addition, in ASLM, to minimize aberrations, the remote focusing objective lens should ideally be identical to the illumination lens, which was easily achievable with this affordable, long-working distance, air objective lens.

Our microscope consists of four main optical components: the laser launch, the ASLM, the illumination arm, and the detection arm (**Fig. 1A-B, Supplementary Figs. 2-3, and Supplementary Table 2**). Briefly, a laser beam with a Gaussian profile enters through the laser launch and is shaped into a light sheet using a cylindrical lens. This light sheet then travels to the ASLM unit, where it is swept by a voice coil and its attached mirror, positioned opposite the ASLM objective lens. The voice coil acts as a single-axis translator, rapidly moving the mirror along the optical axis. Reflecting the primary light sheet through the ASLM and illumination arm results in a shifted light sheet within the chamber. The detection objective lens then collects the fluorescence emitted from the sample, and the image is formed by a tube lens on an sCMOS camera with a rolling shutter (**see online methods**).

**Figure 1.**
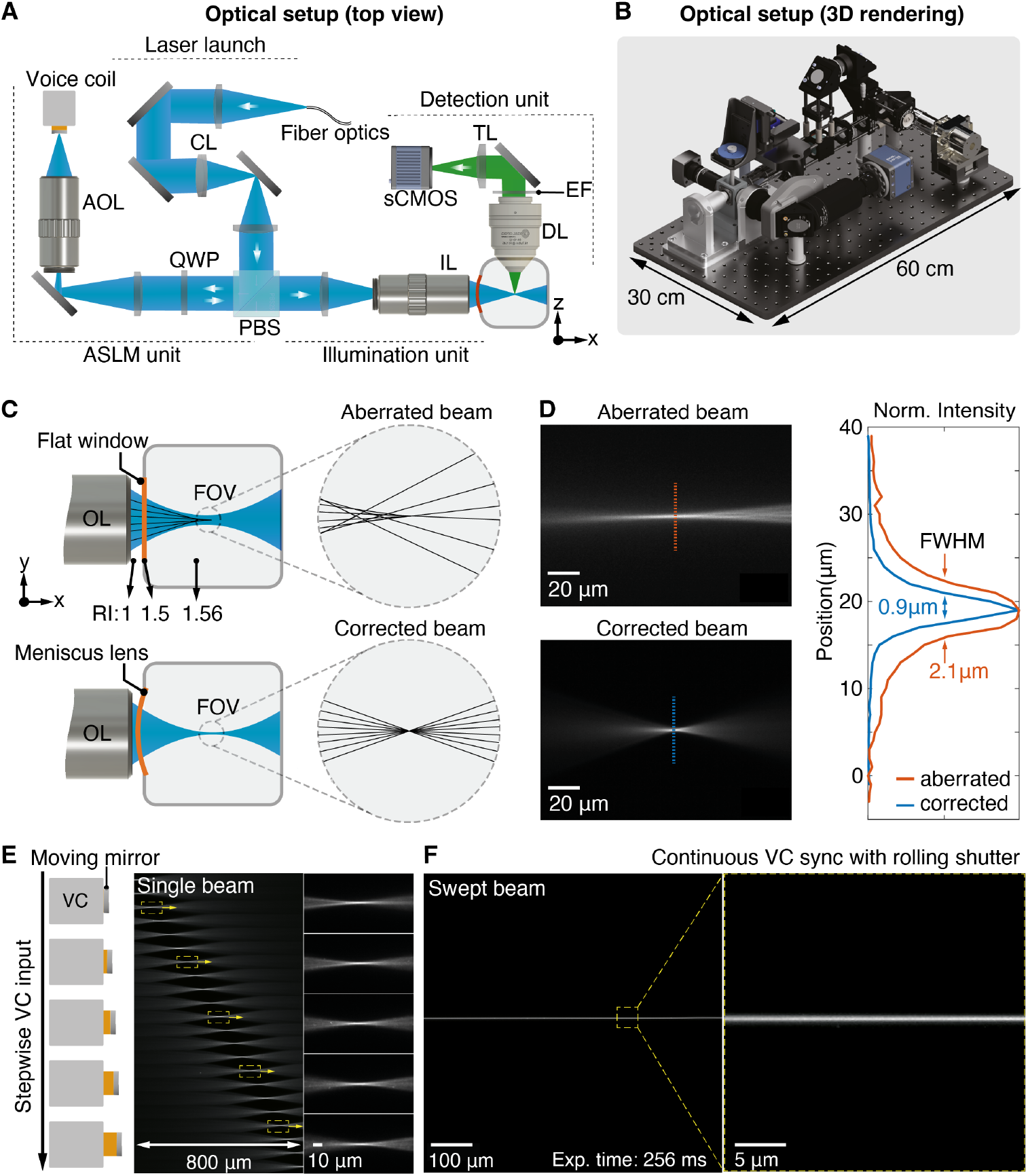
Axially swept cleared tissue light sheet setup and diffraction-limited swept beam. **A**, Optical schematic of the cleared tissue setup from a top view. CL, PBS, QWP, AOL, IL, DL, EF, and TL indicate the cylindrical lens, polarizing beam splitter, quarter-wave plate, ASLM objective lens, illumination objective lens, detection objective lens, emission filter, and tube lens, respectively. **B**, 3D rendering of the optical setup in (A). **C**, Comparison of a glass window and a meniscus lens for correcting spherical aberration in the illumination beam. **D**, single static beam in a chamber with a flat window and a meniscus lens. The graph shows the beam thickness for both cases. **E**, Schematic of the voice coil when its input voltage is changed stepwise. Second and third panels: recorded single beam across the detection field of view, demonstrating the absence of aberrations. **F**, Swept beam with the continuous motion of the voice coil synchronized with the sCMOS rolling shutter.

When using an air objective lens, one of the primary aberrations anticipated is spherical aberration, which prevents the light sheet from achieving the diffraction limit^23^ (**Fig. 1C first row**). To mitigate spherical aberrations, we introduced an off-the-shelf meniscus lens between the air objective lens and the chamber instead of using a flat glass window. The meniscus lens has a curvature that matches the NA of the illumination beam to allow all the rays to enter the chamber perpendicular to the glass surface and to minimize refraction (**Fig. 1C second row**). Comparing the beam quality, we saw strong spherical aberration when using the flat glass window, while it was greatly eliminated with the meniscus lens, reducing the beam size from 2.1 μm to 900 nm, the near diffraction limit (**Fig. 1D**). We also examined the beam quality across the detection FOV to ensure it remained aberration-corrected when sweeping the light sheet. We achieved a homogeneous, confined beam across the entire FOV (800 × 800 μm_2_) by combining ASLM with the meniscus lens. We then synchronized the swept aberration-corrected beam with the sCMOS rolling shutter during the exposure time, recording it over the FOV from left to right (**Fig. 1E-F; Supplementary Movie 2**).

Another essential requirement for ASLM is that the waist of the light sheet forms a straight line from top to bottom across the FOV. This is necessary to synchronize the thinnest part of the sweeping light sheet with the center of the rolling shutter. However, illumination field curvature is expected when using a high NA air objective lens in combination with high RI media across a large FOV (**Fig. 2A**). To evaluate the illumination field curvature, we first directly imaged the light sheet profile using a small mirror inside the chamber at 45 degrees between the optical arms. We then measured the center location of the light sheet cross-section from top to bottom (**see Online methods**). The field curvature of the light sheet was 20 ± 0.2 μm with an excitation beam of 561 nm and RI of 1.56, which was nearly three times larger than the light sheet Rayleigh length (**Fig. 2B, red line**). To correct the light sheet field curvature, we explored replacing the conventional flat mirror in the ASLM unit with a concave mirror with an inverse curvature, as measured by the light sheet field curvature. The chosen off-the-shelf concave mirror (CM127-010-P01, Thorlabs) has a 9.5 mm effective focal length with a maximum offset of 10 μm at 400 μm from its center, which covers the light sheet with an 800 μm height (**Supplementary Fig. 4A**). The field curvature was corrected to less than 1 μm along the light sheet (**Fig. 2B, green line**) compared to the flat mirror. Then, we characterized the axial resolution by imaging 200 nm nanoparticles and measuring the point spread function (PSF). When using the flat mirror, we measured a variety of axial PSFs over the light sheet height from 800 nm to 2.5 μm (**Fig. 2C**). Here, we achieved isotropic resolution only in the central 400 µm of the FOV, while the rest of the FOV remained non-isotropic. However, with the concave mirror, the axial resolution was homogeneous across the entire FOV, with an isotropic resolution of 850 nm. This enabled us to increase the effective FOV by a factor of 2 compared to the flat mirror (**Fig. 2D**).

**Figure 2.**
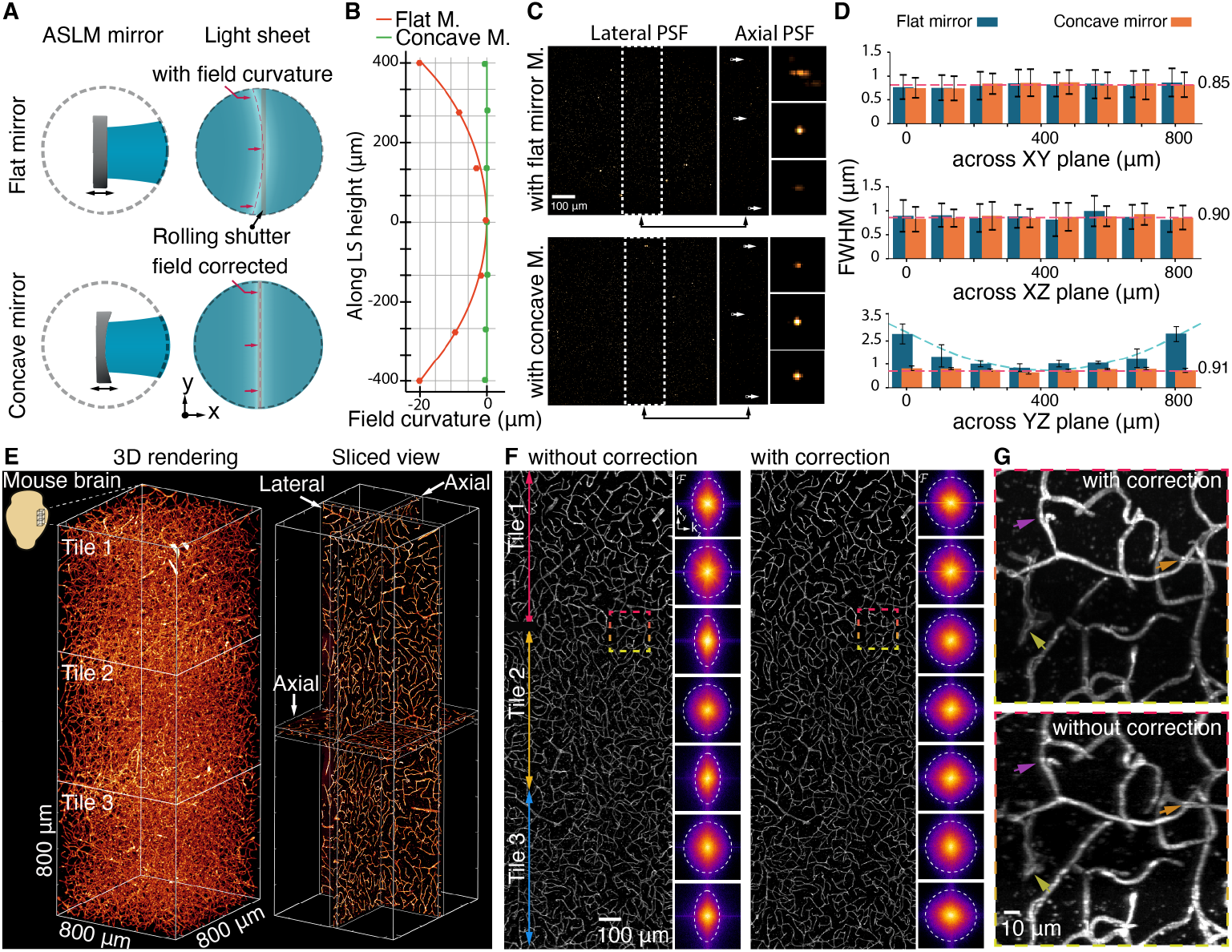
Flat field corrected light sheet enhances FOV and quality across the imaging tiles. **A**, schematic comparison of using the flat mirror and curved mirror in the ASLM unit and their effect on the field curvature of the light sheet. **B**, measured field curvature of the light sheet without (red) and with correction (green). **C-D**, PSF measurement with 200 nm nanoparticles and comparison without and with the field curvature correction in lateral and axial directions. **E**, 3D rendering and middle slices of the lateral and axial views of the recorded data from a cleared mouse brain, stained for micro-vessels with 3 tiles along the height of the light sheet. **F**, axial view of the recorded data in D along the light sheet height and the corresponding Fourier domain comparison with its associated cut of frequency for seven regions. **G**, enlarged view of selected area in D located between adjacent tiles. The arrow heads show the improvements in vessel information with field curvature correction.

To examine the microscope’s performance in cleared tissue, we used a mouse brain labeled with antibodies for brain endothelial cells, visualizing even the smallest vascular structures, so-called capillaries (**Fig. 2E**). The imaging was done for three tiles to evaluate the resolution across the volume, before and after field curvature correction (**Fig. 2F**). The axial view of the recorded images before field curvature correction showed that the cut-off frequency of the non-corrected images varied and decreased to almost half in the Fourier domain compared to the image with corrected field curvature. In contrast, image quality and the frequency distribution remained constant across the tiles when imaged with field curvature correction (**Fig. 2G**). Consequently, eliminating the light sheet field curvature achieved isotropic resolution across a larger FOV and into the corners of the images, crucial for image stitching and quantitative imaging.

Alongside the spatial resolution, the imaging speed is also important for achieving high-throughput imaging of large samples. In ASLM, the maximum speed is mainly determined by the sweeping device, which in our case is a voice coil with a reported speed of up to only 10 Hz for ASLM^13^. To explore the voice coil’s potential to move much faster, we developed a small, closed-loop monitoring arm with a low-power diode laser and a position-sensing detector (PSD, **Fig. 3A**). The axial movement of the voice coil translated into a lateral shift of the reflected laser on the PSD and a modulation of its voltage output (**see Online methods**). Our measurements across typical exposure times from 8 ms to 256 ms confirmed that the voice coil’s response was not uniformly linear across all frequencies, particularly at higher frequencies, which desynchronized the movement of the light sheet and the rolling shutter at several parts of the FOV (**Supplementary Fig. 5**). To eliminate the non-uniformity, a correction waveform was calculated based on the residual of the voice coil’s response to the initial sawtooth waveform, which was then applied back on the voice coil as a correction waveform. Remarkably, even after just a few iteration cycles, the voice coil’s movement became uniformly linear, allowing for precise synchronization between the sweeping light sheet and the sCMOS rolling shutter at almost full speed without compromising FOV coverage (**Fig. 3B-C**). This approach also allowed us to synchronize the voice coil with the rolling shutter at a frame rate of 100 fps (**Supplementary Table 1**).

**Figure 3.**
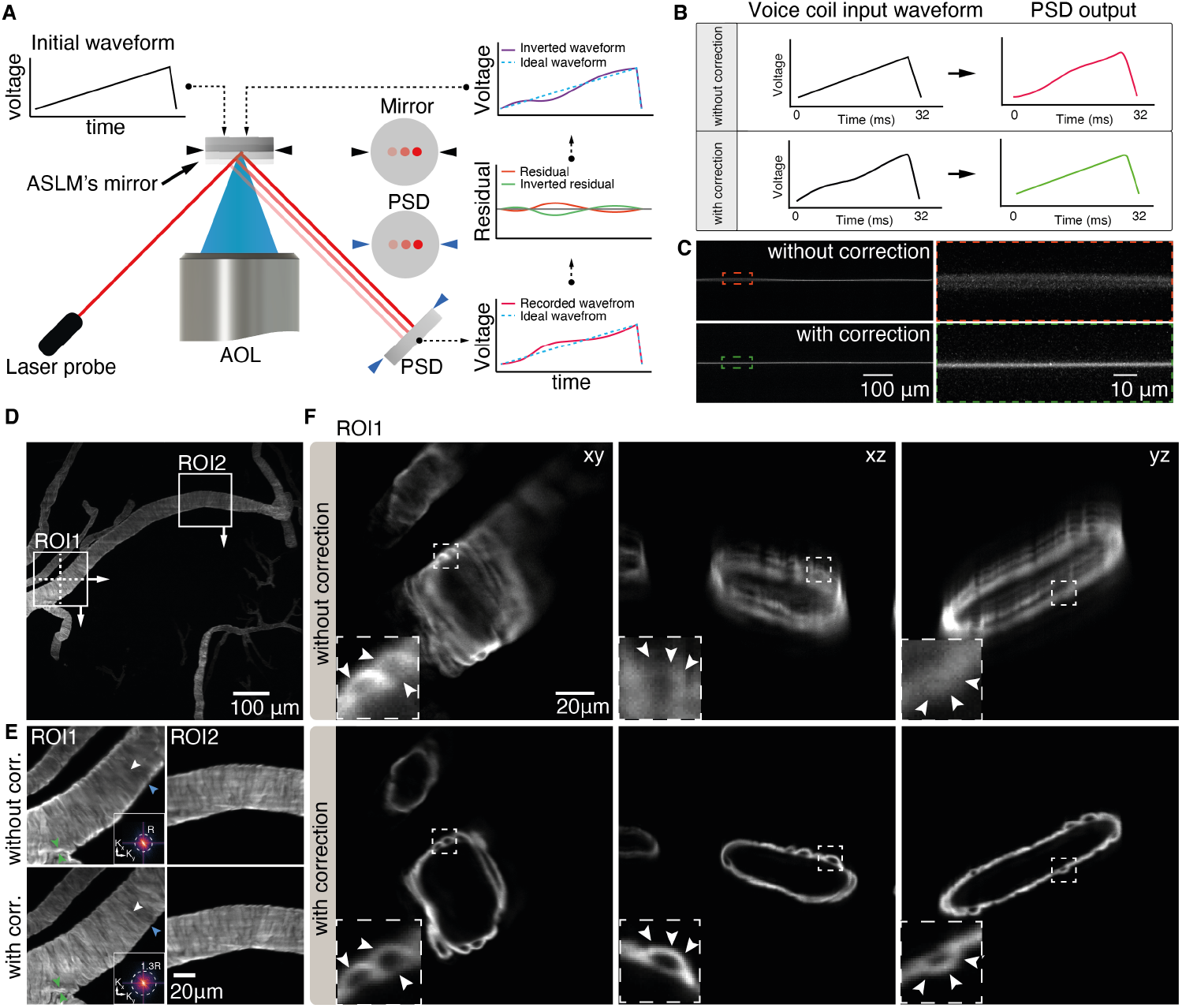
High-speed isotropic resolution achieved by voice coil waveform adaptation. **A**, Schematic of the PSD arrangement and the closed-loop control of the voice coil in the ASLM unit, along with the adaptive algorithm for correcting voice coil movement. PSD indicates the position-sensing device. **B**, Recorded feedback waveform from the PSD without and with correction at voice coil frequency 31 Hz. **C** Recorded swept beam inside the chamber corresponding to the waveforms in B with an exposure time of 25 ms. **D**, Recorded mouse brain stained for smooth muscle actin, showing a selected ROI with and without waveform correction. **E**, Selected ROI1 and ROI2 in D. Resolution analysis in the Fourier domain shows an increase in resolution after voice coil correction. **F**, Cross-section of ROI1 from D in the xy, xz, and yz planes with and without voice coil movement correction. The subsets and arrow heads highlight the same area.

We next tested the system’s performance for high spatiotemporal resolution imaging of a piece of cleared mouse brain (800 × 800 × 2400 μm^3^) with fluorescently labeled larger vessels, especially arteries and arterioles, using an antibody detecting smooth muscle cells (α-smooth muscle actin, αSMA; **Fig. 3D**). Here, the imaging was done with an sCMOS camera with a controllable rolling shutter at its maximum frame rate (PCO Panda; **Supplementary Table 1**). The recorded data showed a resolution variation across the FOV and along the sweeping light sheet in both lateral and axial views before the voice coil waveform correction. Notably, the isotropic resolution across the entire FOV was only achieved with the corrected waveform of the voice coil. Individual fine vasculature structures were resolved in all directions at sub-micron scale at 40 Hz across the entire FOV (**Fig. 3E-F**).

## Benchmarking the light sheet system by imaging cleared specimens at different refractive indices

To benchmark the microscope’s performance for rapid and isotropic imaging of whole cleared organ samples, we first imaged an entire mouse cochlea. The cochlea offers an intricate snail-shaped 3D structure with the spiral ganglion sampling sound information along the frequency axis. iDisco cleared mouse cochlea^10^ (see Online methods) were matched with an RI of 1.56 (ECi, **Fig. 4A**). Immunolabelling with parvalbumin (PV) allows for robust visualization of the entire population of spiral ganglion neurons (SGNs) within the Rosenthal canal. We first imaged a small part of the cochlea to see how much more information can be retrieved along the axial direction when using a thin swept light sheet compared to a conventional light sheet with a thickness of about 5 μm covering the entire FOV: detailed observation, especially in the axial direction, had hitherto remained elusive when using conventional light sheet methods (**Fig. 4B-D**). The typical volume of a cochlea is approximately 1.5 × 1.5 × 2 mm^3^ (**Fig. 4E-F**) and could be imaged in four tiles. We employed a plane spacing of 380 nm, consistent with the pixel size, for an RI of 1.56 to achieve a resolution of 850 nm. Up to 4,000 imaging planes were needed in each tile to cover the entire volume. Because of the isotropic resolution, each SGN was clearly distinguished from its neighbors, and neurites were detected (**Fig. 4G-I**), enabling us to segment each SGN properly (**Fig. 4J-K**). Alongside sub-micron isotropic resolution, our microscope allowed us to image a mouse cochlea with four tiles in just 2.1 to 8.5 minutes at an exposure time of just 8 to 32 ms, respectively (**Supplementary Movie 3**).

**Figure 4.**
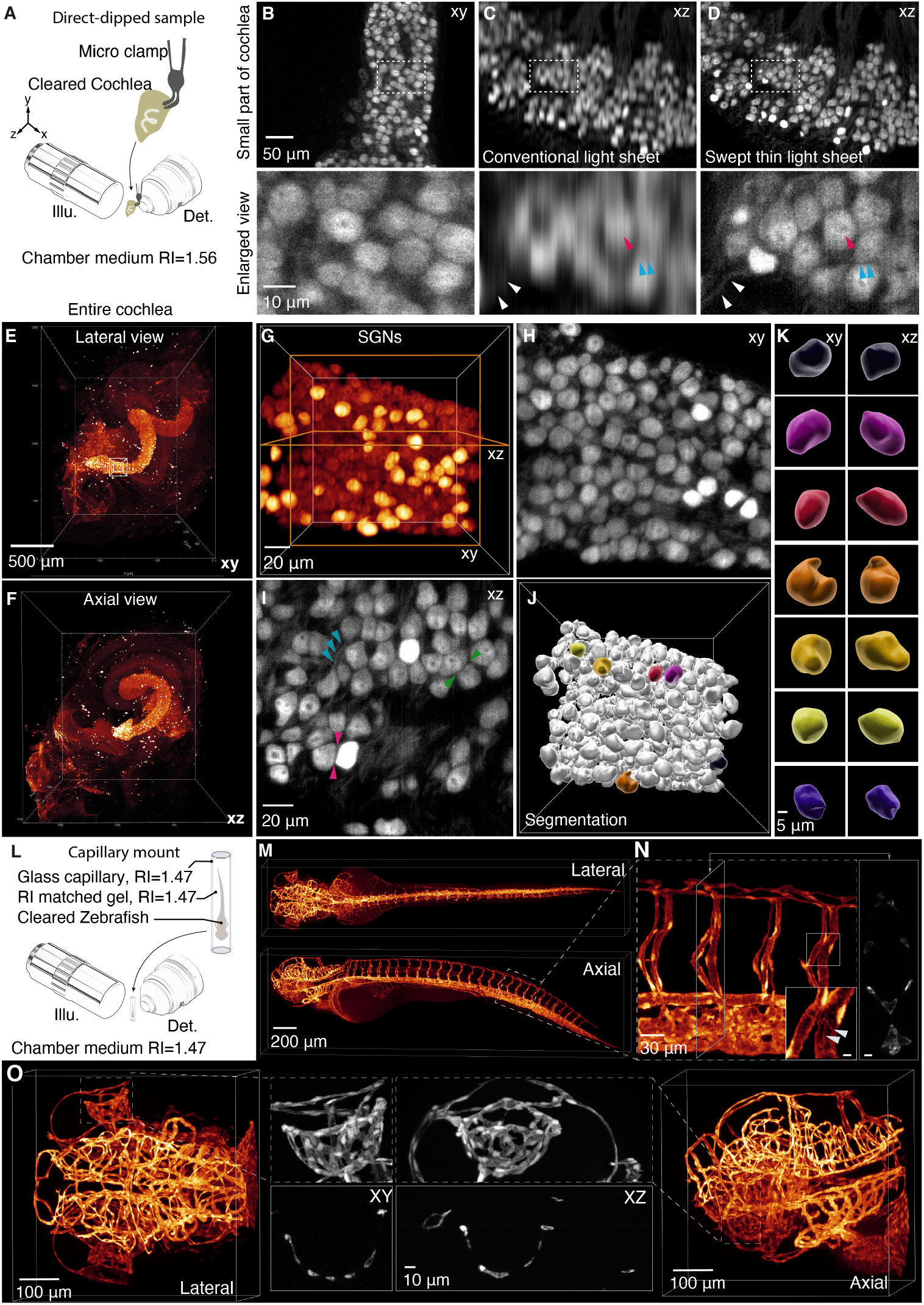
Imaging of a mouse cochlea with isotropic resolution. **A**, schematic of the cochlea and its mounting in the microscope. **B**, Top: recorded data from a small section of an anti-parvalbumin-stained cochlea with a swept thin light sheet. Bottom: selected and magnified region of interest (ROI) from the top row. **C**, Top: reconstructed axial image in B recorded with a conventional light sheet covering the entire FOV. Bottom: selected and magnified ROI from the top row. **D**, Top: reconstructed axial image in B recorded with a swept thin light sheet. Bottom: selected and magnified ROI from the top row. All arrows of the same color indicate comparisons between the magnified images in C and D. **E**, 3D reconstruction of a mouse cochlea from 4 tiles (lateral view). **F**, 3D reconstruction of E (axial view). **G**, a region of interest in the cochlea base with spiral ganglion neurons (SGNs) from E. **H–I**, lateral and axial view of the mid-plane in G. Blue arrows mark exemplary neurites of the bipolar SGNs. The green and red arrowheads indicate the well-resolved gap between the neurons due to the isotropic resolution. **J**, 3D rendering after segmentation in G. **K**, selected segmented SGNs in J (lateral and axial view), labeled with a unique color for each SGN. **L**, schematic of the zebrafish and its mounting in the microscope using a borosilicate glass capillary. **M**, 3D reconstruction of the zebrafish from both lateral and axial views. **N**, enlarged view of a selected region of interest (ROI) in the tail; the inset highlights magnified vessels within this ROI (scale bar: 5 μm). All arrows indicate the endothelial cell boundaries. The right panel displays a cross-section of the tail (scale bar: 10 μm). **O**, enlarged views of the zebrafish head from M, including lateral and axial views (first and last). The middle images represent maximum intensity projections (MIPs) of the vasculature around the eye in both orientations, along with a central cross-section.

To evaluate the microscope’s performance at different refractive indices, we cleared a 3 dpf Tg(kdrl:GFP) zebrafish using an adjusted version of the EZ Clear protocol^20^ (see Online Methods). The zebrafish was mounted in an RI-matched agarose gel inside a borosilicate glass capillary suspended in glycerol (**Fig. 4L**). This mounting method allows for easy mounting of delicate or fragile samples while maintaining a uniform RI (see Online Methods). The sample was imaged in four tiles, totaling 8000 planes for the entire sample, with a plane spacing of 390 nm. The entire sample is recorded with 100 frames per second within an acquisition time of 1 minute and 20 seconds, (**Fig. 4M-O**). The data showed consistent image quality in both lateral and axial views (**Fig. 4M, O**), with fine details such as endothelial cell boundaries clearly resolved in all directions (**Fig. 4N inset**). Isotropic resolution was confirmed by examining the delicate ophthalmic vasculature in both lateral and axial views, which showed equally well-resolved vascular walls and lumina (**Fig. 4O and Supplementary Movie 4**).

### Isotropic 3D imaging enables analysis of cochlear connectomics

To demonstrate the biological applicability and resolution performance of our light sheet microscope in structured and functionally critical tissue, we performed high-resolution imaging of cochlear afferent innervation. Using triple-color fluorescent labeling, we visualized inner hair cells (IHCs), SGNs, and their peripheral neurites (**Fig. 5A-B**). Imaging was carried out at the system’s maximum isotropic resolution, enabling a comprehensive 3D reconstruction of cochlear architecture with consistent detail across all spatial orientations. We first rendered whole-organ 3D overviews with clearly separated channels, followed by the selection of a region of interest (ROI) from the basal turn encompassing SGNs and their neurites (**Fig. 5C-E and Supplementary Movie 5**). Within this ROI, we extracted three cross-sectional planes perpendicular to the local neurite trajectory at distinct locations along the cochlear spiral. In these planes, individual neurites could be resolved and counted, indicating that fine anatomical structures remained sharply defined regardless of orientation. This underscores the system’s isotropic resolution and its suitability for neuroanatomical quantification.

**Figure 5.**
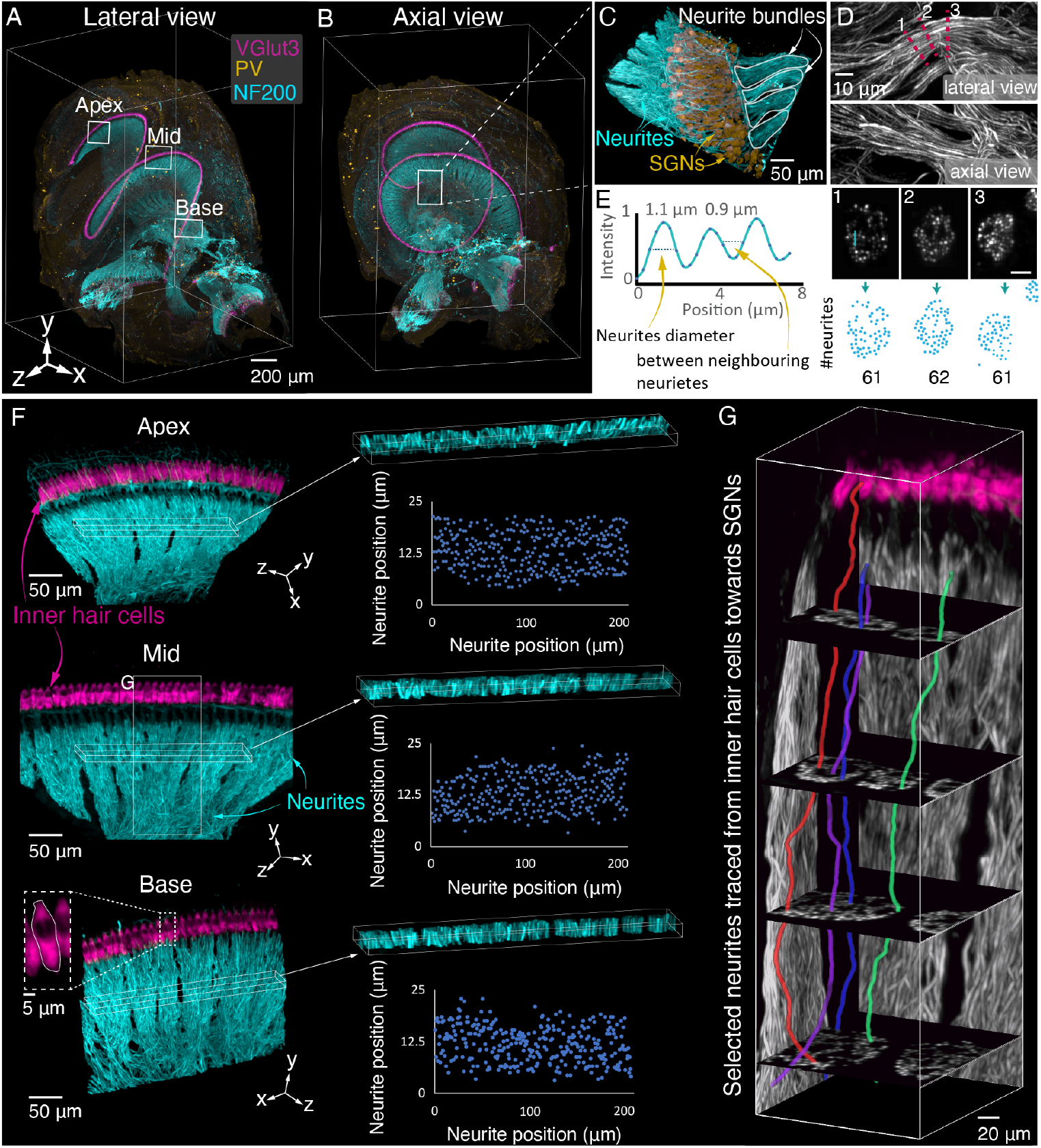
Multicolor imaging of the mouse cochlea. **A, B**, lateral and axial views of a full cochlea volume showing inner hair cells (IHCs) immunolabelled for vesicular glutamate transporter 3 (VGlut3), spiral ganglion neurons (SGNs) labelled with anti-parvalbumin (PV), and neurites labelled with anti-neurofilament 200 (NF200) in x, y, z. **C**, selected and magnified region of interest (ROI) from B, showing neurites and SGNs (left) and neurites in lateral and axial views (right). **D**, analysis of a single neurite bundle in C. First row: lateral view; second row: axial view; third row: neurite diameter and position in three oblique planes along the neurite extension marked with a red dashed line and numbered 1, 2, 3 (scale bar: 10 μm); fourth row: neurite count from exemplary cross-sections shown in D, first row. **E**, line profile of three neighboring neurites in cross-section 3 in D. **F**, selected ROIs from the apical, middle, and basal turns of the cochlea, as shown in A, depicting IHCs labelled with VGlut3 and neurites labelled with NF200 to visualize neurite positions for exemplary counting across all cochlear turns. **G**, tracing SGN neurites from the organ of Corti to the spiral ganglion in an ROI from the mid-turn of the cochlea, shown in D.

To expand the analysis, we selected three larger ROIs from the apical, middle, and basal turns. In each, we quantified the number of SGN neurites terminating on 25 adjacent IHCs—a direct proxy for IHC–SGN connectivity, which is critical for understanding auditory function and pathology (**Fig. 5F**). The ability to perform these counts across the cochlea consistently reflects the uniform image quality, enabling comparative assessments of innervation density and spatial patterning along the tonotopic axis.

We then traced and segmented selected neurites from their synaptic connection with IHCs to the corresponding SGN soma within the Rosenthal’s canal, capturing their full 3D trajectories (**Fig. 5G**). This level of structural detail supports studies of cochlear neuronal connectivity, allowing reconstruction and analysis of afferent pathways at the single-cell level. Such detailed mapping opens the door to systematic studies of cochlear connectomics, including afferent synaptic organization of type I and II SGNs, molecularly defined type I SGN subtypes ^29–31^, efferent connectivity, and circuit-level remodeling in health and disease.

### Imaging of the whole mouse brain

To assess the system’s performance in imaging even larger specimens, we imaged an entire mouse brain (14 mm x 10 mm x 7 mm) immunolabelled for arteries and arterioles by αSMA-targeting antibodies (**Supplementary Fig. 6 and see Online methods**). A compact preview system with a 0.3x optical magnification was installed opposite the detection lens, providing a large overview of approximately 15 mm x 15 mm (**Fig. 6A**). The preview assists the user in choosing the desired orientation and selecting multiple tiles across the entire sample, each covering a full detection FOV of 800 μm x 800 μm.

**Figure 6.**
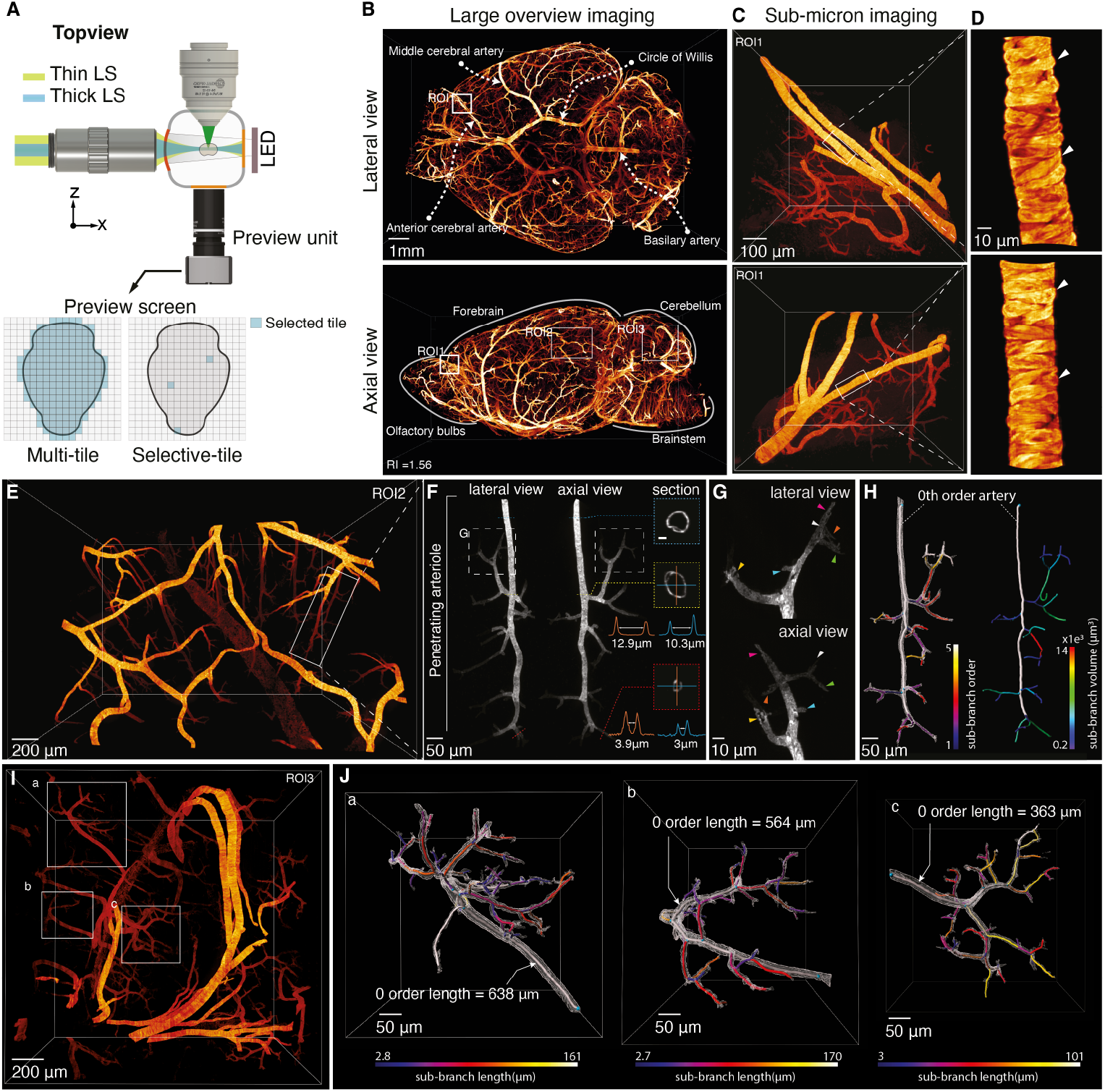
Hybrid imaging of an entire cleared mouse brain. **A**, schematic of the setup with chamber, lenses, and a small camera unit for capturing a preview image of a large sample and selecting the tiles. **B**, entire reconstructed mouse brain from lateral and axial views. **C**, re-imaged selected ROI in B and the reconstructed axial view (second row). **D**, enlarged view of selected small vessel in C (first row) and reconstructed axial view (second row), highlighting individual vessel cells (arrow heads). **E**, selected ROI2 from the cortex in B, which was recorded with the highest possible isotropic resolution. **F**, selected penetrating arterier and cross-sections at different depths and branches(scale bar: 5 μm). The vertical line (orange solid line) and horizontal line (blue solid line) profiles indicate the measured line profiles from the cross-sections of the 0-order artery and the last SMA-positive termini. The inner distance between the artery walls was measured and indicated with a white double arrowhead line. **G**, enlarged view of one of the penetrating sub-arteries in F. The color-coded arrowheads indicate the last SMA-positive arterial termini with the same color from both lateral and axial views. **H**, left: segmentation of the entire penetrating artery and the counted sub-branches until the last SMA-positive termini. Right, the calculated volume of the main and sub-arteries. **I**, selected ROI3 from the cerebellum in B, which was recorded with the highest possible isotropic resolution. **J**, full segmentation of three selected penetrating arteries named a, b, and c. The segmentation and artery tracing enabled us to find the length of the main and sub-arteries in all directions.

We first acquired a quick overview of the mouse brain by selecting all the tiles covering the brain and imaging them with a thick light sheet (3.8 μm). This reduction in axial resolution is quickly achieved by limiting the NA of the illumination beam with an adjustable slit. To achieve an isotropic overview, the pixel size of the recorded image was then matched to the light sheet thickness by a binning of 10. The entire mouse brain was recorded and reconstructed at this resolution in less than 30 minutes. The reconstructed data enabled us to identify the entire brain’s vasculature trees, including the main arteries located at the surface as well as deeper arterioles (**Fig. 6B**). To observe the vessels’ sub-cellular structure, we identified regions of interest (ROI), e.g., part of the anterior cerebral artery, and re-imaged the related selected tile volume (800 × 800 × 800 μm^3^) again at the highest possible isotropic resolution (850 nm), which only took about 21 seconds for 2100 layers (**Fig. 6C**). The raw data provided isotropic subcellular information of individual smooth muscle cells viewed from any angle without deconvolution or other processing (**Fig. 6D and Supplementary Movie 6**).

To further benchmark the biological relevance and resolution fidelity of the system, we have imaged two distinct parts of the mouse brain, targeting anatomically and functionally different regions: the cerebral cortex and the cerebellum. Imaging was carried out at the maximum achievable resolution, allowing us to visualize fine vascular structures throughout the entire volume without compromising depth or field of view. In each region, we isolated and analyzed three individual penetrating arteries, each differing in depth and spatial orientation. We were able to accurately measure vessel diameter along their full trajectories, regardless of spatial orientation or local deformation—demonstrating the system’s ability to maintain uniform, isotropic resolution. This overcomes a major limitation of conventional LSFM systems, which often suffer from degraded axial resolution (**Fig. 6E-F**).

Using 3D segmentation and skeletonization, we reconstructed the full branching architecture of each vessel down to the last SMA-positive termini (**Fig. 6G**). Precise illustration of the arteriolar-capillary transition zone allowed for clear distinction of different types of mural cells without additional markers or tissue cutting. The ring-like morphology of smooth muscle cells in the arterioles was replaced by an enwrapping morphology in higher-order vessels, reflecting ensheathing pericytes^32^. Interestingly, mural cells in higher-order branches appeared less intense, suggesting a reduced expression of smooth muscle actin in those cells. The full reconstruction enabled precise quantification of branch number, length, and morphology (**Fig. 6H and Supplementary Fig. 7**), supporting applications that rely on high-fidelity structural analysis, such as vascular remodeling, such as angiogenesis during development or vascular remodeling after ischemic insults.

In the cerebellum, vascular features—including the lengths of penetrating arterioles, branching patterns, and compact geometries—were also readily recorded. These characteristics, which differ from those in the cortex and reflect the anatomical and functional distinctions of this brain region, were visualized without the need for angle-specific imaging, highlighting the versatility and consistency of our system (**Fig. 6I-J**).

Overall, this analysis highlights how our microscope enables orientation-independent, subcellular-resolution imaging across large tissue volumes. The ability to accurately trace and quantify individual vessels in 3D, regardless of geometry or orientation, makes it a powerful tool for investigating brain vasculature, microcirculation, and disease-associated structural changes at unprecedented clarity and scale.

## Discussion

High-resolution 3D imaging and quantitative analysis of large biological samples are essential for understanding complex structures, such as neuronal circuits and vascular networks. Maintaining spatial context throughout the entire volumes allows for precise and efficient extraction of morphological and functional insights. To meet these needs, we have developed a rapid, aberration-corrected light-sheet microscope for imaging cleared tissues with isotropic sub-micron resolution. This approach demonstrates that detailed morphometric and connectivity analysis across whole specimens has now become possible without requiring tissue reorientation, flattening, distortion, or sectioning.

The microscope utilizes an axially swept light sheet to achieve isotropic resolution over a large FOV. We found that the light sheet field curvature can be effectively corrected by replacing the flat mirror in the ASLM with an off-the-shelf concave mirror rather than using an expensive, dedicated objective lens. The straightened light sheet is key to achieving an isotropic point spread function (PSF) across the extended FOV and stitching volume tiles accurately. While we have only corrected the field curvature in our illumination arm, the same principle could be extended to remote focusing in the detection arm for fast volumetric imaging, as sometimes required in live imaging.

Furthermore, we minimized spherical aberrations and achieved a near-diffraction-limited light sheet using identical, easily exchangeable high-NA air objective lenses in both the illumination and ASLM arm. Combined with an off-the-shelf meniscus lens, we achieved sub-micron resolution. Aberrations in the illumination arm are minimized because both objective lenses are identical. Again, this concept represents a cost-effective solution that could also be extended to the detection arm in future developments. The curvature of the meniscus lens is chosen to match our primary NA of 0.4, but can be adapted to other lenses as needed.

Our closed-loop technique for calibrating and monitoring the ASLM unit enables us to achieve the very high frame rate of 100 Hz. The system controls several aspects of operation, such as pre-calibrating the voice coil’s running waveform and monitoring the voice coil’s movement for any drift, e.g., due to environmental factors like temperature. Our technique can adapt to various voice coil frequencies, such as the typical sCMOS with 40, 100, and 122 Hz frame rates in light sheet mode (Supplementary Table 1), making it universally applicable and robust. Further advancements in sCMOS technology with additional light sheet modes and potentially higher frame rates can be incorporated without sacrificing FOV. Furthermore, our approach may apply to other microscopy techniques that rely on precise remote focusing^33^.

The striping artifacts typically associated with light sheet microscopy are less significant when samples are well-cleared and RI-matched as shown. However, residual striping artifacts can be mitigated using multi-directional selective plane illumination microscopy (mSPIM)^34^ or computational methods^35^. The mSPIM could be seamlessly integrated into our microscope by employing a fast resonance mirror or an acousto-optic deflector (AOD) with kHz-level scanning. Nevertheless, some samples may not achieve perfect RI matching due to hard tissues like bone^36^, which may shift the light sheet and detection plane from the perfect focus^37^. We have addressed this issue by axially fine-tuning the light sheet position through voice coil offset adjustments and defocusing the light sheet to the detection plane using a motorized mirror placed after the cylindrical lens. Finding the optimal imaging angle may become important in future developments, potentially leveraging smart rotation^38^. While our proposed method has been applied within a conventional light sheet configuration with two objective lenses, all techniques introduced in this study can be readily adapted to various other light sheet configurations with different numbers of lenses for diverse applications, including live tissue imaging.

In summary, our cleared tissue microscope is a robust, compact system built from commercially available, off-the-shelf components. We achieved isotropic, sub-micron resolution in all directions with high-speed imaging for high-throughput imaging of biological samples. Our light sheet microscope can reveal the complex sub-cellular architecture across large intact tissues, providing new insights for biological and medical research.

## Online methods

### Optical setup

The optical setup consists of six units: laser launch, remote focusing, illumination arm, chamber, detection arm, and preview unit.

1. **Laser launch:** The setup can either be used with a fiber-beam laser engine (CLE laser engine, 20 mW each laser line, TOPTICA) or a free-beam laser engine (OBISCellX, 100 mW each laser line, Coherent). Each laser engine provides four laser lines at 405, 488, 561, and 640 nm. When using a fiber-coupled laser, the laser beam is collimated with a 25 mm lens, which gives a 7 mm beam. For the free laser beam, a 50 mm lens is paired with a 25 mm lens to provide a 2X beam expansion, which also provides a beam with a 7 mm diameter at the 561 nm laser line. Following the beam collimation and expansion, an initial light sheet is formed with a cylindrical lens with f = 45 mm. This primary light sheet is subsequently collected by a secondary lens with f = 60 mm. A polarizing beam splitter (PBS) that guides the beam through the ASLM arm is positioned in front of this secondary lens.
2. **Remote focusing**: The beam polarization is controlled by rotating the optical fiber to achieve maximum reflection at the PBS towards the ASLM. After passing the PBS, the laser beam encounters a sequence of optical elements: a quarter-wave plate (QWP), a 100 mm lens, a mirror, and a 20x objective lens before hitting the ASLM mirror. The quarter-wave plate converts the laser beam’s linear polarization into circular polarization (e.g., clockwise). After reflection from the ASLM mirror, the beam returns with a circular polarization in the opposite direction (counterclockwise). When the beam passes through the QWP again, it regains linear polarization but in an orthogonal direction, enabling it to pass through the PBS and proceed to the illumination objective lens. The ASLM unit has a voice coil actuator (Thorlabs Blink) to sweep the light sheet. This setup includes either a flat mirror with a diameter of 12.5 mm (Thorlabs) or a concave mirror (Thorlabs CM127-010-P01) with a 9.5 mm focal length, which corrects the light sheet field curvature while simultaneously sweeping the light sheet.
3. **Illumination arm:** Following the PBS towards the main illumination arm, a 60 mm focal length lens collects the reflected beam after the remote focusing. The lens back focal plane is conjugated to the PBS center, while its front focal plane aligns with the illumination objective lens back focal plane (Mitutoyo 20X NA 0.4, MY20X-804). After the illumination lens, the beam encounters a positive meniscus lens with a 100 mm focal length (Thorlabs LE1234). This lens acts as a correction plate, preventing spherical aberrations when transitioning from air to a medium with a higher RI in the chamber.
4. **Chamber:** The chamber is made out of aluminum with four openings. One is used for holding the meniscus lens (Thorlabs LE1234). Two of them hold two glass windows with a thickness of 1 mm (Thorlabs WG10110), one for the preview camera and one for the preview LED. The last opening is used for the dipping detection objective lens. All the glass and lenses are sealed with sealant O-rings.
5. **Detection unit:** The detection unit uses a multi-immersion dipping objective lens (ASI multi-immersion lens 16X NA 0.4, 15-10-12) with a working distance of 12 mm at RI 1.45. The emission light is filtered through one of the six emission filters mounted on a custom-made filter wheel equipped with a robotic servomotor. For standard operation, the image is formed by a 200 mm tube lens (TTL200-A, Thorlabs) onto an sCMOS chip with a rolling shutter (PCO Panda 4.2m or Hamamatsu Fusion C14440-20UP) featuring a 6.5 μm pixel size and a 2048×2048 pixel square chip. Alternatively, a 160 mm tube lens was used for an sCMOS chip with a 4.6 μm pixel size (PCO 10bi CLHS).
6. **Stage and sample holder:** For volumetric imaging, the specimen is continuously moved through the light sheet in the Z-direction. This process captures a block tile using a linear stage (PI M112) at plane spacings of 0.38 μm, mirroring the detection system pixel size, when the RI is at 1.56. Here, we used tiles in the X- and Y-dimensions for specimens that exceed the detection system’s FOV, using another two linear stages (PI M112).

### Refractive Index compatibility

The meniscus lens (positive meniscus, focal length 100 mm) was selected to match the curvature of the illumination beam’s wavefront, which is determined by the numerical aperture of the illumination objective. This design ensured that the beam enters the sample chamber from air into higher RI media at a nearly perpendicular angle across the entire beam profile. Consequently, the performance of the meniscus lens remained unaffected by the RI of the chamber medium, making it compatible with all common clearing protocols. Similarly, the concave mirror used in the ASLM path compensated for the field curvature introduced by the overall optical setup of the illumination arm. Its function is also independent of the chamber RI. Therefore, switching between different immersion media requires no replacement of the meniscus lens or the concave mirror.

### Compensation of chromatic aberrations for multi-wavelength imaging

We used an air objective lens in the illumination arm, which, in combination with the high refractive index of the clearing solvent, can introduce chromatic aberrations. This resulted in wavelength-dependent shifts in the position of the light sheet along the sweeping direction. To correct for this shift and ensure that all wavelengths focus at the same position, we experimentally determined the optimal offset voltage for the voice coil actuator at each excitation wavelength. As we performed multi-color imaging sequentially, we loaded a separate, pre-calibrated waveform for each laser wavelength, each with the appropriate offset voltage applied. This ensured that light sheets of different wavelengths are spatially aligned with the rolling shutter, enabling accurate multi-color imaging without chromatic aberration.

### Data streaming, post-processing, and visualization

The microscope is operated by a compact computer (NUC 11) running on an Ubuntu operating system. The data collected by the sCMOS is seamlessly streamed from the NUC to an external solid-state drive (Samsung T7, 4TB) via a Thunderbolt connection during standard operations, maintaining a frame rate of under 40 fps. For higher frame rates exceeding 40 fps, a 2TB M2 (Samsung NVMe) drive housed in an enclosure that supports Thunderbolt 4 is used, allowing for a streaming speed of 40 Gbps. Each imaged volume tile is directly stacked in BigTiff or RAW format as it is written to the external SSD. Subsequently, the data is converted into HDF5 format to enable lossless compression for Fiji ^2^ and BigStitcher ^40^ for further analysis and image stitching. All 3D rendering and segmentation have been done using Imaris 9.7.

### Point spread function measurement

The microscope’s point spread function (PSF) was measured by imaging 200 nm gold nanoparticles embedded in clear, RI-matched agarose gel. First, 1.5% hard-gelling agarose is prepared with 0.2 percent gold nanoparticles already vortexed for about 5 minutes to avoid clustering. It is mixed with the prepared agarose gel and formed with a 2 mm glass capillary in a cylinder shape for easy holding. Once solidified, the gel is pushed out with a plunger into ethanol for dehydration. The dehydration process is done in three steps using 20, 50, and 100 percent ethanol for 15 minutes, 15 minutes, and 30 minutes, respectively. After dehydration, the agarose gel is transferred to the final RI-matching solvent for at least one day. Once RI-matched, the agarose is transferred to the same capillary or glued to a sample holder for PSF measurement. For the PSF measurement, we imaged the nanoparticles using a 561 nm laser line with a power of 0.5 mW, a z-step size of 380 nm, and a 20-pixel camera rolling shutter. All parameters corresponded to the same slit size and z-step as used for imaging the biological samples. The PSF is then determined based on the full width at half maximum (FWHM) of the intensity profiles of the recorded nanoparticles using Fiji^39^.

### Closed-loop monitoring arm with a position sensing device

We have used a position sensing device (PSD, PDP90A, Thorlabs) with a K-Cube controller (KPA101, Thorlabs) in combination with a compact probe laser module with a Gaussian intensity profile (1 mW laser, 635 nm, PL202, Thorlabs). The PSD and probe laser are positioned at 45 degrees relative to the optical axis of the ASLM unit. The laser is reflected on the ASLM mirror and hits the PSD. The axial movement (along the optical axis) of the voice coil translates into the lateral movement of the laser spot on the PSD. By tracking the center position of the laser beam on the PSD, the output voltage of the K-Cube controller (Output X) generates a corresponding signal with an accuracy of 0.7 μm at a wavelength of 635 nm (approximately 100 μW at the PSD). To achieve this level of accuracy, we have used an absorptive neutral density (ND) filter with 10 % transmission (NE510A, Ø1/2”, Thorlabs) attached to the PSD. It should be noted that the closed-loop monitoring is used only during calibration to optimize the voice coil waveform for the desired scan frequency. During actual imaging, the system operates with the pre-calibrated waveform.

The output of the K-Cube is connected to a compact multi-function generator (Moku:Go, Liquid Instruments), which streams the PSD output directly to the controlling computer for further analysis (**Supplementary Fig. 8**). The signal is then processed using a Python script that calculates the residual of the PSD output compared to the initial linear waveform and generates a corrective waveform. The corrective signal is sent to the voice coil via a function generator. The system operates using a feedback closed-loop algorithm, which works by either converging to a predefined metric or incorporating user feedback based on the swept beam quality (**Supplementary Note 4)**. The best corrective waveform is selected, typically achieved in fewer than 10 iterations. Ultimately, the optimal corrective waveform for each target frequency needs to be calculated only once.

### Field curvature measurement

To measure the light sheet field curvature, we used the PSF measurement and direct imaging of the light sheet cross-section. The PSF measurement was the same as described. However, for the light sheet cross-section measurement, we used a small mirror of 2 mm x 2 mm glued to a glass capillary. The mirror is placed in the chamber between the objective lenses at 45 degrees, directly reflecting the light sheet into the detection arm. Without any field curvature, the cross-section of the light sheet should be a straight line from top to bottom of the field of view. However, in the presence of field curvature, the light sheet thickness varies along the FOV height. To measure the field curvature, we first parked the light sheet in the center of the detection FOV (the voice coil was parked at the center position with zero amplitude). Then, the light sheet cross-section is recorded in several positions by gently moving the mirror along the FOV. The intensity profile of the light sheet for recorded cross-sections is measured from top to bottom. Then, the thinnest light sheet in each plane along the FOV height is selected. The relation between these thinnest parts of the light sheet describes the field curvature.

### Animals

Animal handling complied with relevant national and international guidelines for the care and use of laboratory animals in research (European Guideline for Animal Experiments 2010/63/EU) and in accordance with regulations put forward by the state authorities of Lower Saxony (LAVES) and Schleswig-Holstein (MLLEV). Mice were kept in a 12-hour light/dark cycle with ad libitum access to food, water and enrichment. Mice euthanized for extracting cochleae and brains in this study aided the development of the new microscope presented here, capable of high-throughput and high-resolution sub-micron imaging to further address various biological questions in following studies. Adult zebrafish (*Danio rerio*), embryos and larvae were maintained following established protocols. The transgenic line *Tg(kdrl:GFP)* was used, where GFP is expressed in the vascular endothelium. Zebrafish embryos were kept at 28.0 ° in 0.3X Danieau’s medium with 0.003% 1-phenyl-2-thiourea (PTU) added at 1 day post fertilization (dpf) to suppress pigmentation. Embryos were checked for potential malformations and expression of the endogenous reporter using a fluorescence stereomicroscope. Larval zebrafish at 3 dpf showing the desired expression pattern and without apparent malformations were selected for sample preparation.

### Sample preparation (Mouse Cochlea)

After extraction from the temporal bone, cochleae were fixed for 45 minutes in 4 % formaldehyde, subsequently rinsed in phosphate-buffered saline (PBS) and decalcified for up to one week in 10 % ethylenediaminetetraacetic acid (pH 8). During decalcification, cochleae were continuously trimmed with surgical instruments (Fine Science Tools) using a bright field microscope (Zeiss; Olympus SZ61) to remove as much bone as possible for easy access and later penetration of the light sheet beam during optical sectioning.

### Immunolabelling and tissue clearing (Mouse Cochlea)

For robust and reproducible immunolabelling as well as optical clearing, we used a newly adapted version of the optimized iDISCO^+^ clearing protocol adapted to the cochlea (cDisco^+^)^10^. Here, we first decalcified the mouse cochlea for 4 days using 10 % ethylenediaminetetraacetic acid of pH 8.0 at room temperature (RT). Second, we decolorized the decalcified specimen for up to 3 days at 37 °C using 25 % amino alcohol (N,N,N’,N’-Tetrakis(2-Hydroxypropyl)ethylenediamine). Next, we employed pretreatment and permeabilization of the samples for approximately 4 days according to iDisco^19^, specifically without the use of methanol, followed by epitope blocking at 37 °C. For reliable immunolabelling of the entire population of spiral ganglion neurons within the Rosenthal canal, sufficient penetration of the cochlea was achieved by incubating each antibody for 14 days at 37 °C followed by washing in PBS-Tween 0.2 % with Heparin 10 μg/mL for up to 1 day^10^. Here, we used a guinea pig polyclonal anti-parvalbumin antibody (Synaptic Systems, 195 004) at a dilution of 1:300, pre-conjugated with a camelid single domain anti-guinea pig antibody labelled with Atto565 (NanoTag Biotechnologies, N0602) at a molar ratio of 1:3. Further, we used a monoclonal mouse anti-neurofilament 200 antibody (Sigma-Aldrich, N5389) at a dilution of 1:400 and a polyclonal rabbit anti-vesicular glutamate transporter 3 antibody (Synaptic Systems, 135 203) at a dilution of 1:300. For a secondary staining staining an anti-mouse antibody labelled with Alexa 647 (Invitrogen, A-21236) and an anti-rabbit antibody labelled with Alexa 488 (Invitrogen, A-11008), each at a dilution of 1:200 were used. Following the immunolabelling, the cochleae were dehydrated, stepwise, in a methanol/H^2^O series ranging from 20 to 100 % methanol for up to 1 day and then delipidized with dichloromethane at RT^41^. Finally, RI-matching was performed by initial embedding in dibenzyl ether for 1 day and consecutive transfer to ECi for storage and imaging at RT.

### Sample mounting (Mouse Cochlea)

For imaging and sufficient matching of the RI the cleared cochleae remained in ECi for at least 6 days. The cochleae were then carefully mounted on a customized 3D-printed sample holder. Here, we clamped the cochlea dorsal of its basal turn at the attached vestibular system to the holder and inserted the mounted cochleae with the cochlear apex facing the bottom of the imaging chamber. Thus, the modiolus of the cochlea is placed approximately perpendicular to the applied light sheet beam within the imaging chamber.

### Mouse brain

The whole procedure of preparing, staining, and clearing the tissue was performed according to a published protocol^42^ with slight adaptations. The method is based on the iDISCO procedure and optimized to visualize the brain vasculature. Briefly, after decapitation, the samples were extracted by opening the skull and removing the brain, ideally including the meninges and surface vessels. In addition to perfusion fixation using 4 % paraformaldehyde in PBS, a post-fixation period of 3 h at RT followed the brain extraction. The tissue was dehydrated using ascending methanol concentrations and incubated in 100 % methanol for 2 h at RT. Lipids were extracted by incubation in dichloromethane/methanol (2:1) overnight at RT and samples were bleached using 5 % H2O2 in methanol overnight at 4 °C. Rehydration was performed by descending methanol concentrations and samples were kept for 24 h at 37 °C in PBS / 0.01 % sodium azide. Blocking was performed using PBS supplemented with Triton-X 0.2 % and porcine gelatin 0.2 % for 24 h at 37 °C. Subsequent incubation with primary antibodies lasted for 10 days at 37 °C in the blocking solution. Here, primary antibodies included antibodies detecting smooth muscle cells by using anti-α smooth muscle actin (αSMA) directly coupled to Cy3 (Millipore, C6198, 3.75 µg/ml) and antibodies detecting endothelial cells by using anti-podocalyxin (R&D systems, MAB1556, 0.5 µg/ml) and anti-CD31 (R&D systems, AF3628, 0.133 µg/ml). After washing overnight, secondary antibodies were then incubated for another 7 days at 37 °C. For comprehensive staining of all endothelial cells in the brain, both primary antibodies were combined and visualized by secondary antibodies carrying the same fluorophore. We used anti-goat and anti-rat secondary antibodies coupled to Alexa647 (Invitrogen, A21447,2.5 µg/ml and Abcam, ab150155, 2.5 µg/ml, respectively). SMA staining did not include a secondary antibody since the primary antibody is directly coupled to Cy3. After staining, the brains were cleared starting with another dehydration and washing in a dichloromethane/methanol mixture as described above. Benzyl ether was used to optically clear and then transfer to ECi for storage and imaging at RT.

### Tissue clearing and sample mounting (zebrafish)

Hatched *Tg(kdrl:GFP)* zebrafish at 3dpf were anaesthetized using 0.02 % MS222 and fixed in 4 % paraformaldehyde in PBS overnight followed by several washes in PBS. For clearing of the zebrafish larvae, an adjusted version of the EZ Clear protocol was used^20^. The specimens were delipidated in EZ Clear solution (50% tetrahydrofuran in Milli-Q water) for 2 hours at RT, followed by washes in Milli-Q. RI matching was performed with an adjusted EZ View solution. As reported in the original protocol (Hsu et al 2022), the RI of the EZ View solution can be tuned by using different concentrations of urea. Starting at 1.2 M urea, the refractive index of the EZ View solution was monitored using a digital refractometer (Krüss DR301-95; A.Krüss Optronic GmbH, Hamburg, Germany) while small amounts of urea were added and dissolved until an RI of n_D_=1.473 was reached (if the RI is too high, it can be decreased by adding 0.2X PBS). This RI was chosen because it allowed for the convenient mounting of delicate samples and immersion as follows: Samples were transferred to a gel consisting of 1.5% low-gelling point agarose in the RI-tuned EZ View and drawn into borosilicate glass capillaries (n_D_ = 1.473; ID = 1.0 mm, OD = 1.5 mm). The RI of the gel was confirmed before mounting and adjusted by adding Milli-Q if necessary. After gelation of the EZ View agarose, the capillaries were sealed at the bottom with plasticine. Because of the identical RI, glycerol was used as immersion medium as it is inexpensive and does not dry out to form a sticky film and is easier to clean than EZ View.

## Acknowledgments

We want to thank Christof Schmidt and his team at the central mechanical workshop of the Department of Physics at the University of Göttingen for their invaluable assistance in creating mechanical components. We also acknowledge Kevin Adner for several electronic components, Paul Maier for his constructive input on the microscope development, and Tabea Quilitz for her feedback on imaging mouse cochlea. We thank Reto Fiolka and Kevin Dean for their insightful discussions on selecting the voice coil. We thank Peter Lenart, Mohammad Amin Eskandari, and Roman Tsukanov for providing the Hamamatsu camera and frame grabber. We thank Beate Lembrich and Zoela Syeda Gilani, both from Lübeck, as well as Ina Preuss and Sina Langer of the Göttingen InnerEarLab for their expert technical support and Bettina Wolf and Jakob Neef for scientific discussion. We further acknowledge funding by the Alexander von Humboldt Foundation (JH; AvH professorship) and the German Centre for Cardiovascular Research (DZHK; 81×2300301) on behalf of MA, JW, and JH. MA, GFM, JH, and TM are supported by the MWK (Niedersächsisches Ministerium für Wissenschaft und Kultur). We further acknowledge the support by the German Research Foundation via the Cluster of Excellence Multiscale Bioimaging (MBExC; EXC 2067/1-390729940) and its Hertha Sponer College at the University of Göttingen (MA, GFM, LR, AMD, TM, and JH) as well as the collaborative research center 889 (LR). The work was additionally funded by the Else Kröner Fresenius Foundation through the Else Kröner Fresenius Center for Optogenetic Therapies (to JH and TM). The mouse brain experiments were supported by a grant from the Deutsche Forschungsgemeinschaft to JW (WE 6456/1-1).

## Author contributions

Designed research: MA, JH. Designed and built the optical setup: MA. Developed preview tiling Python script: MA. Developed TCP/IP pipeline for controlling microscope remotely: GFM, JL. Wrote microscope control system software: JL. Evaluated possible light sheet microscope configuration: MA, KW, JH. Prepared zebrafish: TG. Prepared Mouse Cochlea: LR, AMD. Prepared Mouse brains: JW. Optimised sample mounting: MA, TG. Performed imaging and post-processing: MA, with input from TG, JW, and JH. Performed image analysis: MA, with input from TG, JW, TM, and JH. Provided feedback on microscope performance: TG, LR, AMD, JW, TM, JH. Provided resources for Mouse cochlea experiment: TM. Provided resources for Mouse brain experiment: JW, MS. Provided resources for zebrafish experiment: JH. Wrote paper: MA, JH, with input from all authors. Directed and provided resources for the entire research: JH.

## Data availability

Due to the large size of the imaging datasets in this study, they are not stored in a public repository but can be obtained from the authors upon request.

## Code availability

The scripts are available from the authors upon request. The CAD models of the costumed parts are publicly available via the following link: https://zenodo.org/records/15767193

