## Supplementary for "Isotropic, aberration-corrected light sheet microscopy for rapid high-resolution imaging of cleared tissue"

**Supplementary materials**

Supplementary Note 1. Resolution and depth of field of Multi-Immersion Objective

Supplementary Note 2. Vasculature segmentation and tracing

Supplementary Note 3. Practical considerations for mitigating aberrations

Supplementary Note 4. Voice coil calibration procedure

Supplementary Fig. 1. Resolution calculation for multi-immersion objective lens.

Supplementary Fig. 2. Schematic of the cleared tissue light sheet microscope.

Supplementary Fig. 3. 3D rendering of the cleared tissue optical setup.

Supplementary Fig. 4. Mechanical details of the assembly of the voice coil, position sensing device (PSD), and chamber.

Supplementary Fig. 5. Frequency response of the voice coil measured with the position sensing device.

Supplementary Fig.  6. Mouse brain preparation for imaging.

Supplementary Fig. 7. Segmentation and tracing of two selected penetrating arterioles.

Supplementary Fig. 8. Electronics components and their electrical waveforms.

Supplementary Table 1. Comparison between the tested sCMOS cameras.

Supplementary Table 2. Microscope parts list

**Supplementary Note 1. Resolution and depth of field of the multi-immersion objective lens**

The theoretical resolution and imaging depth of field (DOF) are presented in this section for the ASI objective lens with NA 0.4. The calculated parameters are sorted by varying the tube lens and magnification to make it easier to follow the resolution and DOF behavior (**Supplementary Fig. 1**). The tube lenses are chosen from the 160–300 mm focal length range, which provide magnifications of 13–25×. For tube lenses with focal lengths less than 200 mm, the lateral resolution and magnification increase as the focal length of the tube lens approaches 200 mm. Following that, the resolution remains constant as it is limited by the camera's pixel pitch of 6.5µm. The highest resolution achievable with the objective lens is then given by the objective lens' NA. For instance, for a refractive index of 1.56 and a tube lens of f=200 mm or longer, the highest theoretical achievable resolution for the ASI lens with an NA of 0.4 is 740nm. The depth of field (DOF) of the objective lens decreases as the focal length of the tube lenses is increased. This is because the DOF and magnification have an inverse relationship. Thus, increasing the magnification decreases the DOF. As a result, the optimal tube lenses for the ASI lenses have a focal length of 200 mm to achieve the highest resolution possible.

**Supplementary Note 2. Vasculature segmentation and tracing**

Mouse brain vasculature segmentation was performed using a local contrast method in Imaris 9.6.1 to enhance vessel visibility against the background. The analysis was conducted at the original resolution without any downsampling, and no smoothing was applied to preserve the fine structural details of the vasculature. Each segmentation block consisted of a 3D cropped volume centered on an individual penetrating artery, with an average volume of 500 µm × 500 µm × 500 µm. The segmentation was initially executed using the Surface module in Imaris to generate a volumetric representation of the main artery. Smaller segmented particles that did not belong to the arterial tree were excluded—first through thresholding based on segmented volume, and then through manual inspection to eliminate any remaining irrelevant structures. The resulting clean surface segmentation of the arterial tree was then used as a baseline for the Filament Tracer module in Imaris, which was utilized to automatically trace and reconstruct the artery's sub-branches and the surrounding vascular network.

**Supplementary note 3. Practical considerations for mitigating aberrations**

This study focuses on spherical aberrations and field curvature, as they are the primary limitations in high-resolution imaging across large fields. However, asymmetric aberrations—such as astigmatism and coma—can also occur, typically due to potential optical or mechanical misalignments. In our system, these aberrations were not observed after careful alignment. Since coma aberrations may be caused by off-axis beam propagation, in our setup, the most critical point where this can occur is at the entrance to the meniscus lens. If the beam exiting the illumination objective lens is not well aligned with the optical axis of the meniscus lens, coma aberration can be introduced to the light sheet. Additionally, misalignment between the detection objective's focal plane and the meniscus lens can cause the light sheet to enter off-axis, which may also increase the potential of coma aberration. These issues can be avoided by precisely aligning the optical components and carefully designing the chamber to ensure the meniscus lens optical axis aligns with the detection objective lens's focal plane.

Astigmatism is another expected aberration in conventional light sheet imaging, especially when using glass capillaries to mount samples. However, in our system, this aberration was negligible due to the overall imaging principle, which ensures that the refractive index (RI) of the sample matches that of the chamber medium. For rigid samples, the sample can often be directly immersed in the chamber medium, allowing for complete RI matching between the sample and the medium, which reduces the potential for astigmatism aberration. For fragile samples that require more secure mounting in glass capillaries, minimizing astigmatism relies on matching the RI of the cleared sample and the chamber medium to that of the capillary glass.

**Supplementary note 4. Voice coil calibration procedure**

To calibrate the voice coil movement, we performed the following three steps:

**Step 1 – Empirical measurement of light sheet shift.** We first empirically measured the shift of the light sheet for each 1 mV step and observed that each 1 mV corresponds to a 4 µm shift in the chamber. We also determined that the effective light sheet length (twice the Rayleigh range) is 8 µm. To maintain optimal optical thickness, the voice coil movement must remain within 4 µm (i.e., the Rayleigh range). This sets our predefined metric: the maximum residual error in the sweep should not exceed 4 µm, corresponding to 1 mV.

**Step 2 – Iterative waveform correction.** We recorded the response of the voice coil to a linear ramp function using the PSD. The recorded data was smoothed to reduce noise. We identified the minimum and maximum of the smoothed signal and aligned them with the endpoints of the ramp. The residual waveform was calculated by subtracting this signal from the ideal ramp. We quantified both the peak residual and its standard deviation (σ). A sweep was considered acceptable when the peak residual was below 1 mV and σ below 0.24 mV. If these criteria were not met, the inverted residual was added to the input to generate a corrected waveform. This iterative process—measure, correct, reapply—was repeated until both thresholds were met. This ensures the beam remains synchronized with the rolling shutter and within the optical limits imposed by the Rayleigh range.

**Step 3 – User-guided validation of beam quality.** The refined waveform was then used to drive the voice coil, and the resulting swept beam was evaluated manually. As a secondary quality metric, the beam should maintain consistent thickness across the field of view. Users inspected the beam profile at multiple positions by extracting and comparing line profiles. If any fluctuations in beam size were observed, this indicated residual errors persisted. In such cases, Step 2 was repeated for further fine-tuning.

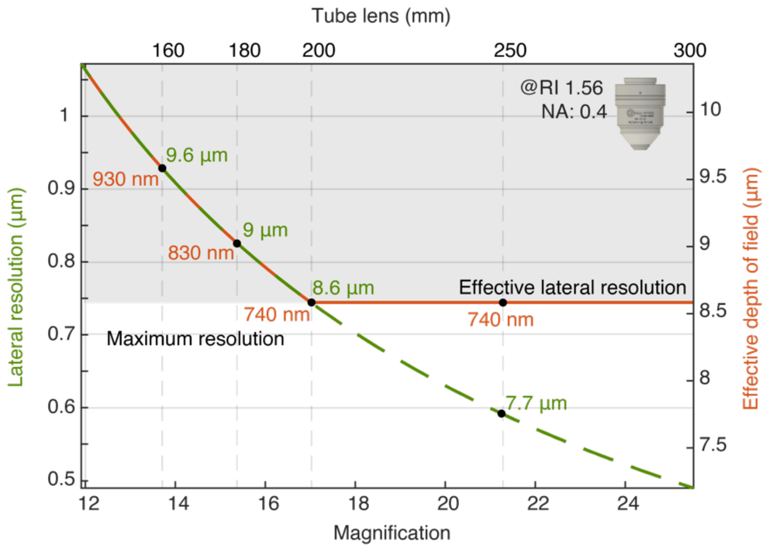


**Supplementary Fig. 1. Resolution considerations for multi-immersion objective lens**. Lateral resolution and effective depth of field for the ASI lens with NA 0.4 are calculated in relation to the magnification and focal length of the tube lens. The solid orange and dashed green lines represent the lateral resolution and DOF, respectively. The grey area indicates the achievable resolution with a camera pixel pith of 6.5µm.

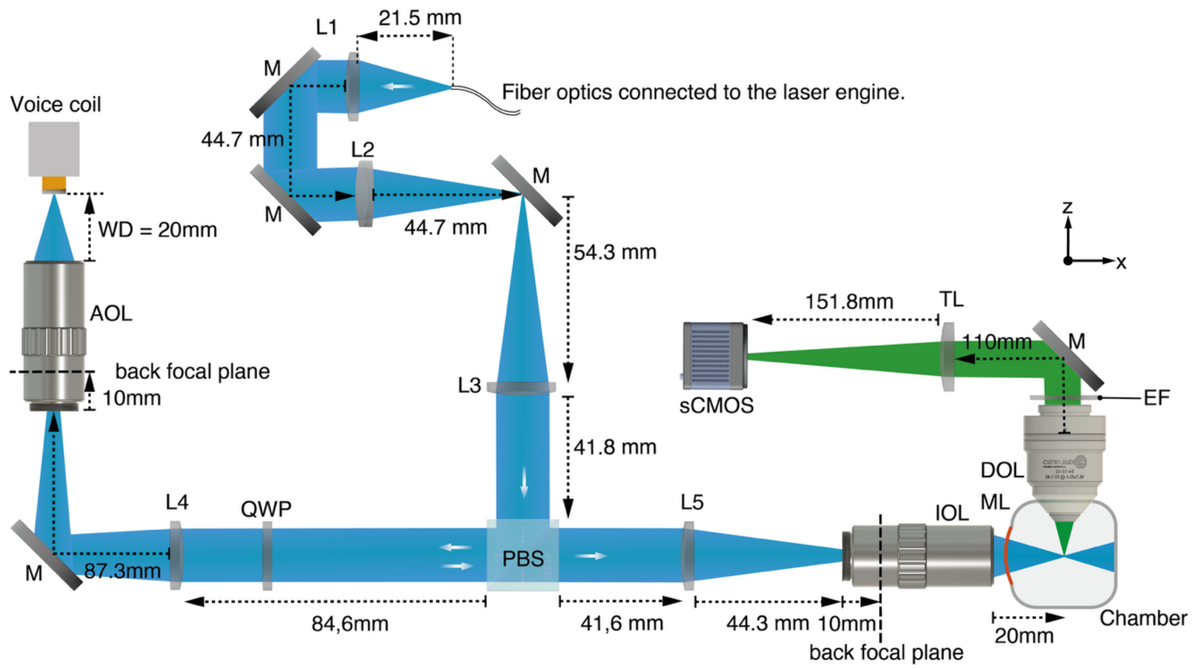


L1: Collimator lens, F = 25 mm, Fb = 21.5 mm (Fb indicates back focal plane)
M: Mirror
L2: Cylindrical lens, F = 50 mm, Fb = 44.7 mm
L3: Expander lens, F = 60 mm, Fb = 54.3 mm
PBS: Polarizing beamsplitter
QWP: Achromatic quarter-waveplate
L4: ASLM's tube lens, F = 100 mm, Fb = 97.3 mm
AOL: ASLM's objective lens, WD = 20 mm, Fb = 10 mm (inside the objective from rear thread)
ASLM's mirror: Concave mirror, F = 9.5 mm
L5: Illumination tube lens, F = 60 mm, Fb = 54.3 mm
IOL: Illumination objective lens, WD = 20 mm, Fb = 10 mm (inside the objective from rear thread)
ML: Positive meniscus lens, F = 100 mm
DOL: Multi-immersion detection objective lens, WD = 12 mm
EF: Emission filter
TL: Detection tube lens, F = 200 mm, WD = 151.8 mm, Pupil distance = 110 mm (working range: 70–170 mm)
sCMOS: Camera with light sheet mode

**Supplementary Fig. 2. Schematic of the cleared tissue light sheet microscope, illustrating the actual physical distances between all optical components.** Dashed arrowheads indicate the precise distance between the faces of two adjacent optical elements. At the bottom of the schematic, abbreviations for each optical element are listed alongside their optical parameters. "F" represents the focal length of the lenses, and "Fb" indicates the back focal plane. All optical components are chosen from achromatically designed optical elements to cover the full visible spectrum.

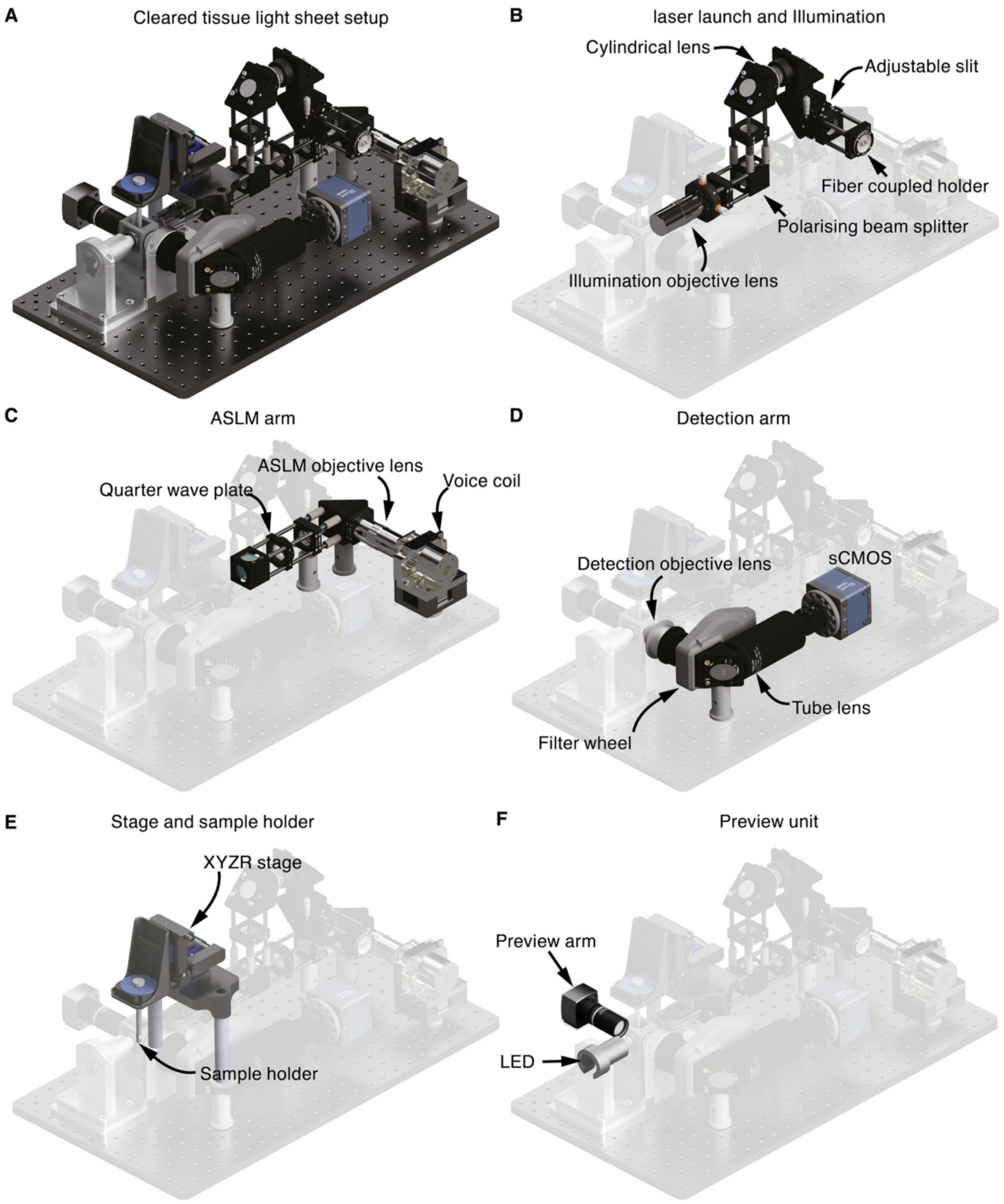


**Supplementary Fig. 3. 3D rendering of the cleared tissue optical setup. A**, 3D rendering of the entire setup. **B**, laser launch and illumination arm. **C**, ASLM arm. **D**, detection arm. **E**, sample stage and holder. **F**, preview unit. The base plate's size is 30x60 $cm^{2.}$

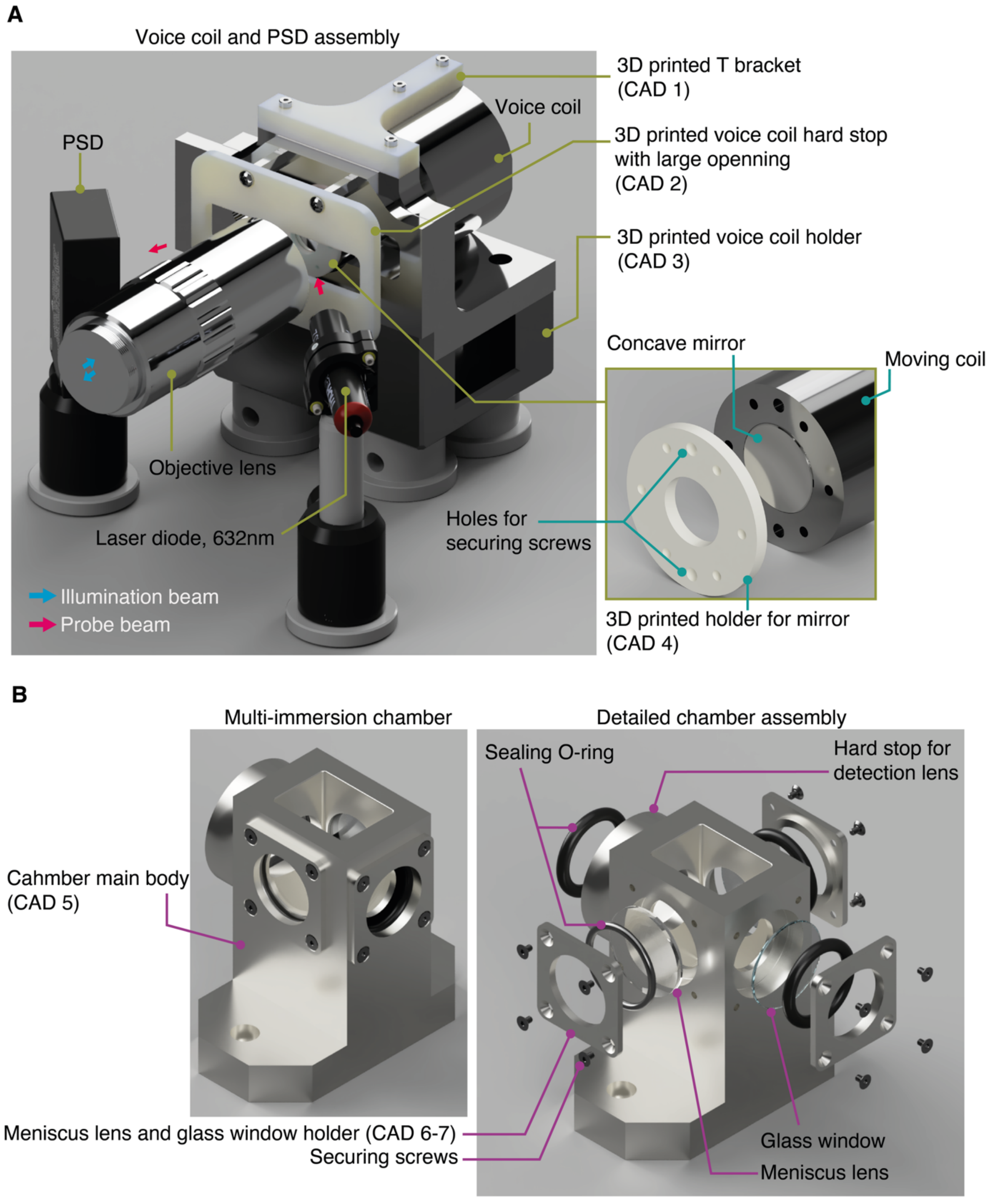


**Supplementary Fig. 4. Mechanical details of the assembly of the voice coil, position sensing device (PSD), and chamber. A**, assembly of the voice coil, including all 3D-printed parts and the holder. A T-bracket is printed and attached to the voice coil to dampen unnecessary vibrations. The hard stop is also 3D-printed using a rigid material, with a widened horizontal opening that allows the probe laser line to enter and reflect seamlessly toward the PSD, as indicated by the red arrowhead. The entire voice coil is mounted and held by a 3D-printed holder, which is supported by stainless steel legs. In the right image, the core of the moving coil is magnified to show where and how the concave mirror is secured using a 3D-printed ring holder. **B**, Left: overview of the CAD model of the multi-immersion chamber. Right: exploded view of all detailed components of the chamber, including holders, meniscus lens, glass windows, and O-rings, as well as the placement of the hard stop for detection. The hard stop is positioned to keep the front of the multi-immersion lens precisely 12 mm (the working distance for all refractive indices) from the illumination optical axis, aligned with the center of the meniscus lens.

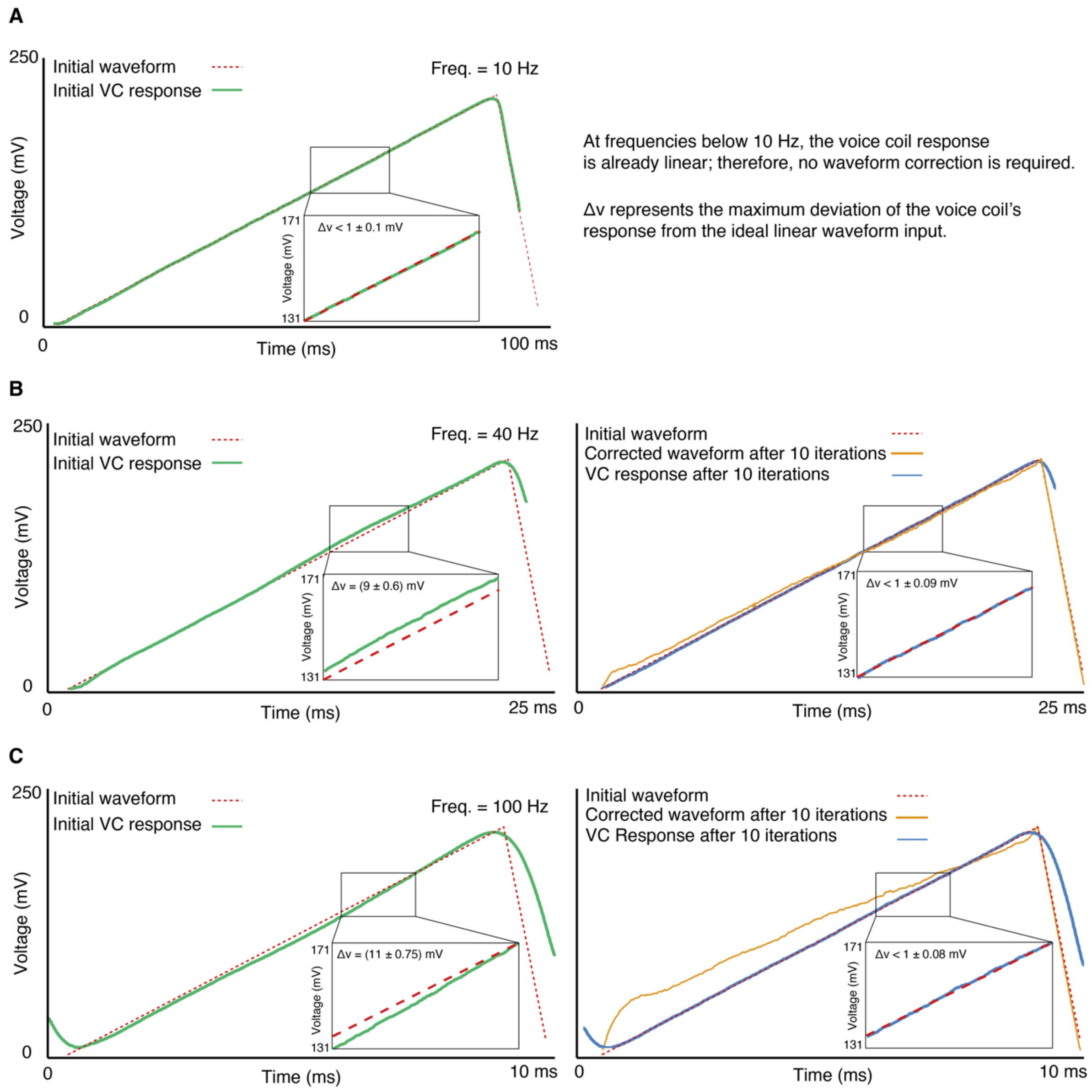


**Supplementary Fig. 5. Frequency response of the voice coil (VC) measured with the position sensing device (PSD). A**, Input ramp waveform at 10 Hz (red dashed line) and the corresponding VC response recorded by the PSD (green solid line). Inset: an enlarged view of the recorded waveform and the calculated maximum deviation. B, Left: 40 Hz ramp waveform and VC response (green solid line). Inset: an enlarged view of the recorded waveform and the calculated maximum deviation. Right: VC response after 10 iterations (green-blue line) and the corrected waveform (orange solid line). Inset: an enlarged view of the recorded waveform and the calculated maximum deviation. **C**, Left: 100 Hz ramp waveform and VC response (green solid line). Inset: an enlarged view of the recorded waveform and the calculated maximum deviation. Right: VC response after 10 iterations (green-blue line) and the corrected waveform (orange solid line). Inset: an enlarged view of the recorded waveform and the calculated maximum deviation.

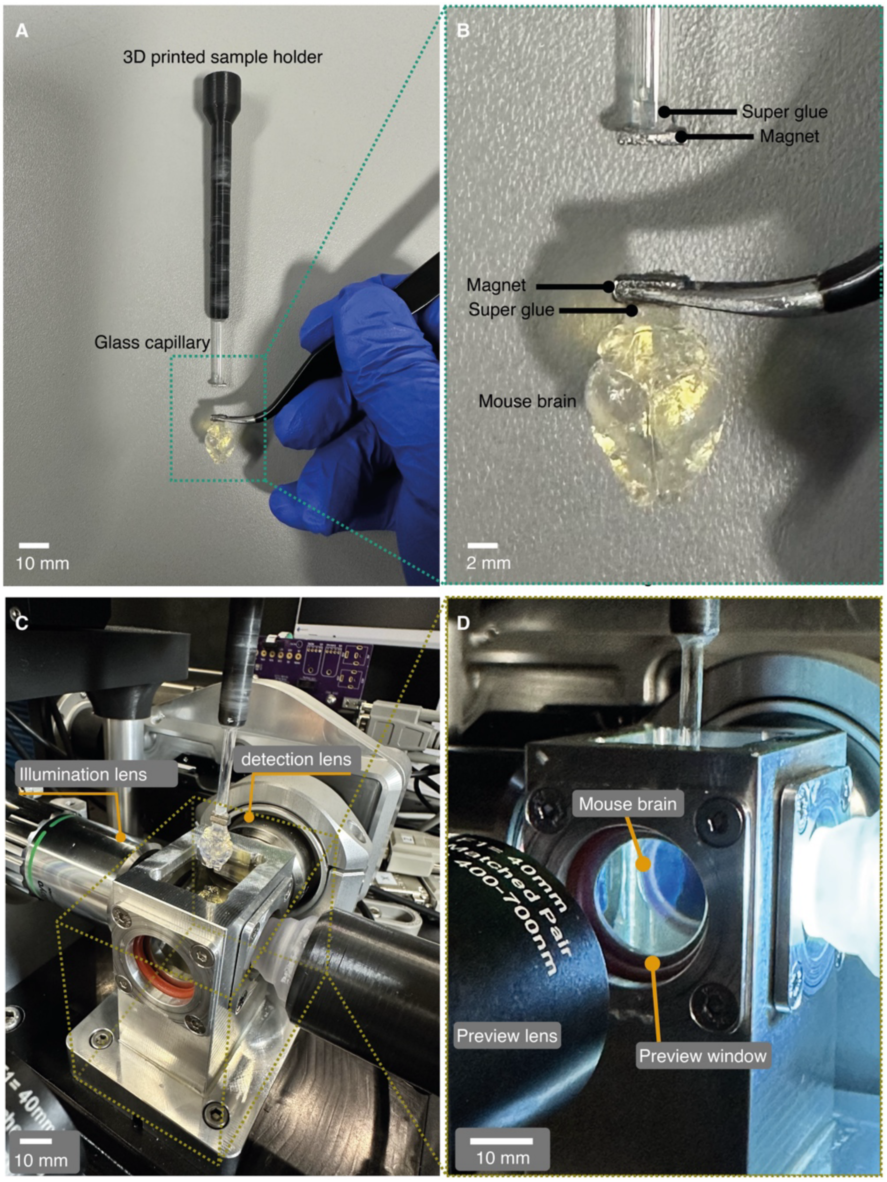


**Supplementary Fig. 6. Mouse brain preparation for imaging. A**, A cleared mouse brain is glued to a magnet using super glue and a 3D-printed sample holder secures a glass capillary with another magnet at its tip. **B**, Enlarged view of the mouse brain shown in A. **C**, An illumination objective lens, a detection objective lens, and an LED with a 3D-printed holder surround a chamber filled with ECi. **D**, The cleared mouse brain inside the chamber is visible through a preview window facing the preview lens.

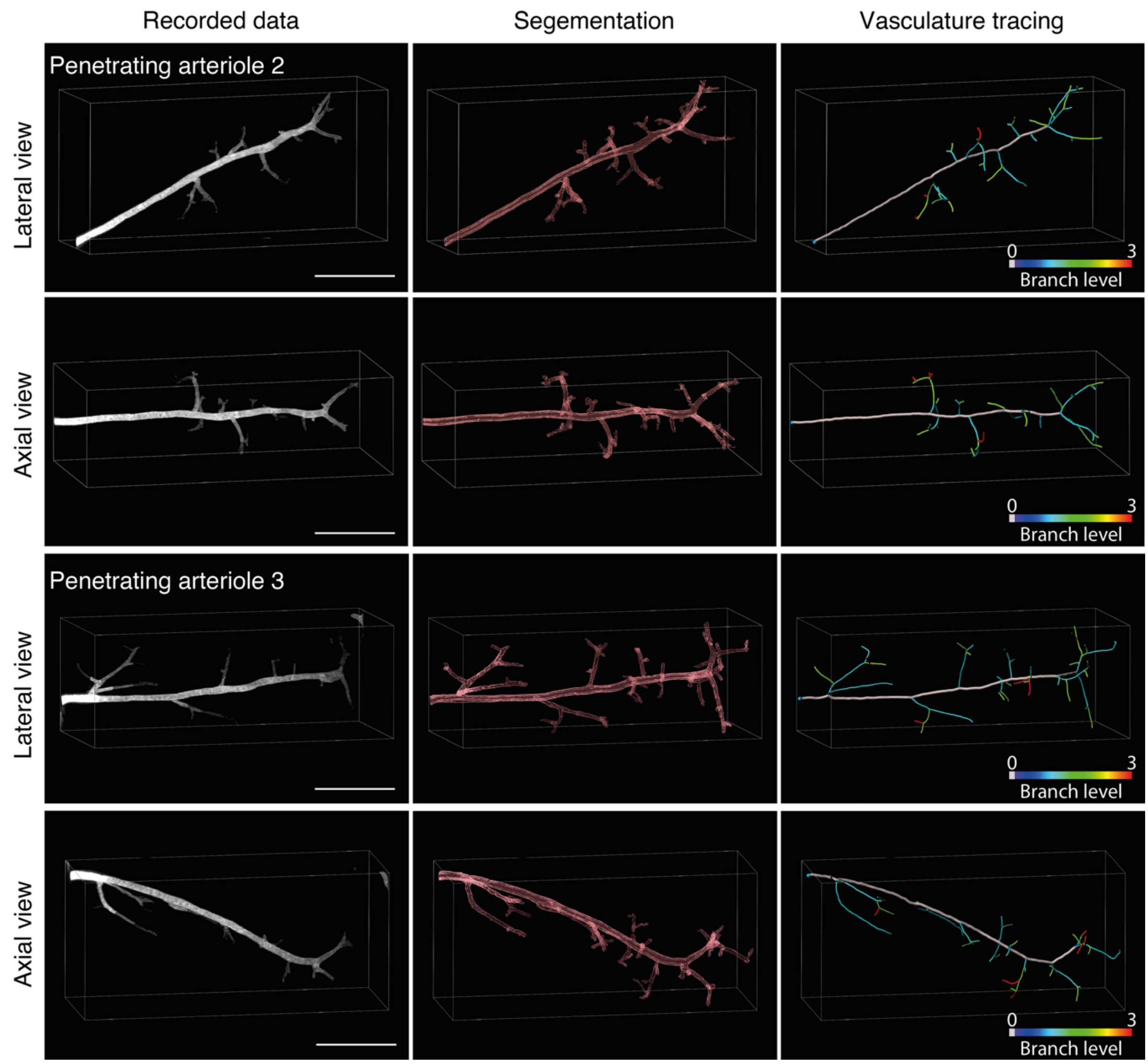


**Supplementary Fig. 7. Segmentation and tracing of two selected penetrating arterioles from Figure 6E.** The first column: the recorded data of two penetrating arterioles in both lateral and axial views. The middle column: segmentation of the corresponding vessels. The third column: vasculature tracing. The color-coded tracing shows the sub-branch levels. Scale bars: 200 µm

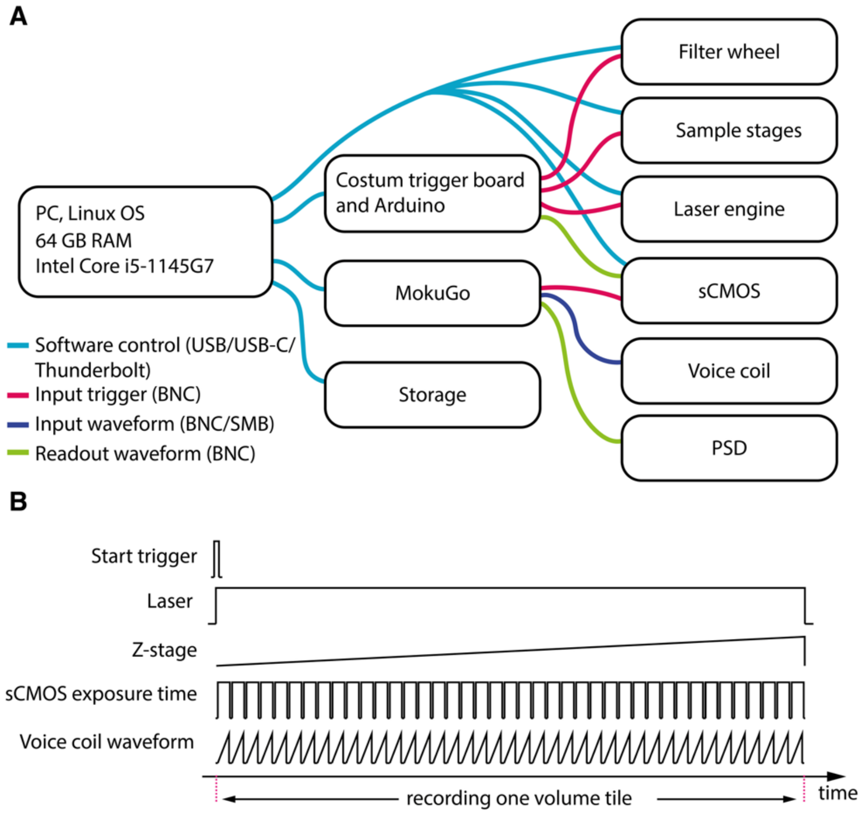


**Supplementary Fig. 8**. **Electronic components and their electrical waveforms. A**, The microscope is run by a small Linux computer. All components are either directly connected to the computer via USB or connected to trigger boards, such as custom boards, Arduino, and MokuGo. The recorded data is streamed to a solid-state drive (SSD). **B**, The microscope's components are started by a user command, and a trigger board sends signals to the other components, such as the laser, z-stage, sCMOS, and voice coil.

**Supplementary table 1. Comparison between the tested sCMOS cameras.**

| sCMOS parameters in light sheet mode | pco.panda 4.2 | Hamamatsu Orca-Fusion | pco.edge 10bi |
|---|---|---|---|
| Maximum frame rate (Hz) | 40 | 100 | 120 |
| Achieved frame rate in ASLM mode (Hz) | 32 | 84 | 100 |
| Exposure line (µs) | 12.136 | 4.868 | 6.84 |
| Chip size (pixel) | 2048x2048 | 2304x2304 | 4416x2368 |
| Used chip (pixel) | 2048x2048 | 2048x2048 | 2368x2368 |
| Effective FOV (µm x µm) | 780x780 (TL: 200mm) | 780x780 (TL: 200mm) | 625x625 (TL: 200mm) 750X750 (TL:160mm) |
| Volume imaging speed of $1mm^3$ (4000 images) | 2.2 min | 45 sec | 32 sec |
| Camera connection | USB-C | Frame grabber | Frame grabber |

**Supplementary table 2. Microscope parts list**

| | | | | |
|---|---|---|---|---|
| Positive Meniscus Lens | LE1234 | 1 | Thorlabs | 25 |
| Concave mirror | CM127-010-P01 | 1 | Thorlabs | 46 |
| Achromatin lens f: 25 mm | AC127-025-A-ML | 1 | Thorlabs | 100 |
| Achromatin lens f: 60 mm | AC254-060-A | 2 | Thorlabs | 190 |
| Achromatin lens f: 100 mm | AC254-100-A | 1 | Thorlabs | 93 |
| Glass window | WG11010 | 2 | Thorlabs | 187 |
| Adjustable slit | VA100CP/M | 1 | Thorlabs | 328 |
| Polarizing Beamsplitting | CCM1-PBS251/M | 1 | Thorlabs | 376 |
| Emission filter | Filters | 6 | Thorlabs | 2281 |
| Cylindrical lens | ACY254-050-A | 1 | Thorlabs | 478 |
| Tube lens | TTL200-A | 1 | Thorlabs | 573 |
| Mirror | BB1-E02-10 | 1 | Thorlabs | 782 |
| Halfwave plate | 10RP52-1B | 1 | Newport | 1222 |
| Quarterwave plate | 10RP54-3B | 1 | Newport | 1222 |
| Illumination Objective lens | MY20X-804 20X Plan Apochromat | 1 | Mitutoyo / Thorlabs | 2613 |
| ASLM Objective lens | MY20X-804 20X Plan Apochromat | 1 | Mitutoyo / Thorlabs | 2613 |
| Detection objective lens | Multi-immersion Objective lens, 54-10-12 | 1 | Special optics / ASI | 17000 |
| Function generator | MokuGo | 1 | Liquid instrument | 600 |
| Voice coil | BLINK high-speed focuser | 1 | Thorlabs | 5226 |
| Sample stage | 3X M1-112 DC motor, 1X U-628.03 | 1 | Physik instrumente | 18214 |
| Camera | PCO Panda 4.2m | 1 | Excelitas | 8000 |
| Mechanical hardware and 3D prints | Mechanical hardware and 3D prints | 1 | - | 5000 |
| Laser engine | Laser Engine (CLE 25mW) / OBIS CellX 100 mW | 1 | Toptica / Coherent | 27700 |
| | | | **Total cost** | **94844** |
